# Duplication and divergence of the retrovirus restriction gene *Fv1* in *Mus caroli* mice allows protection from multiple retroviruses

**DOI:** 10.1101/802363

**Authors:** Melvyn W. Yap, George R. Young, Renata Varnaite, Serge Morand, Jonathan P. Stoye

**Affiliations:** The Francis Crick Institute, London, UK; Centre National de la Recherche Scientifique - Centre de coopération Internationale en Recherche Agronomique pour le Développement ASTRE, Faculty of Veterinary Technology, Kasetsart University, Bangkok, Thailand; Faculty of Medicine, Imperial College London, London, UK

**Author notes:** Corresponding author: (JPS). Current address: Centre for Infectious Medicine, Karolinska Institute, Stockholm, Sweden (Renata Varnaite). These authors contributed equally to the work (Melvyn W. Yap, George R. Young).

## Abstract

Viruses and their hosts are locked in an evolutionary race where resistance to infection is acquired by the hosts while viruses develop strategies to circumvent these host defenses. Forming one arm of the host defense armory are cell autonomous restriction factors like Fv1. Originally described as protecting laboratory mice from infection by murine leukemia virus (MLV), Fv1s from some wild mice have also been found to restrict non-MLV retroviruses, suggesting an important role in the protection against viruses in nature. To begin to understand how restriction factors evolve, we surveyed the *Fv1* genes of wild mice trapped in Thailand and characterized their restriction activities against a panel of retroviruses. An extra copy of the *Fv1* gene, named Fv7, was found on chromosome 6 of three closely related Asian species of mice (*Mus caroli*, *M. cervicolor* and *M. cookii*). The presence of flanking repeats suggested it arose by LINE-mediated retrotransposition. A high degree of natural variation was observed in both *Fv1* and *Fv7*, including numerous single nucleotide polymorphisms resulting in altered amino acids, as well as insertions and deletions that changed the length of the reading frames. These genes exhibited a range of restriction phenotypes with activities directed against feline foamy virus (FFV), equine infectious anemia virus (EIAV) and MLV. It seems likely, at least in the case of *M. caroli*, that the observed gene duplication confers protection against multiple viruses not possible with a single restriction factor. We suggest that EIAV-, FFV- and MLV-like viruses are endemic within these populations, driving the evolution of the *Fv1* and *Fv7* genes.

**Author Summary:** During the passage of time all vertebrates will be exposed to infection by a variety of different kinds of virus. To meet this threat, a variety of genes for natural resistance to viral infection have evolved. The prototype of such so-called restriction factors is encoded by the mouse *Fv1* gene, which acts to block the life cycle of retroviruses at a stage between virus entry into the cell and integration of the viral genetic material into the nuclear DNA. We have studied the evolution of this gene in certain species of wild mice from South East Asia and describe an example where a duplication of the *Fv1* gene has taken place. The two copies of the gene, initially identical, have evolved separately allowing the development of resistance to two rather different kinds of retroviruses, lentiviruses and spumaviruses. Independent selection for resistance to these two kinds of retrovirus suggests that such mice are repeatedly exposed to never-before-reported pathogenic retroviruses of these genera.

## Introduction

Retroviruses are obligate parasites that usurp the host machinery for propagation, inserting their genomes within those of their hosts as an integral part of their life cycles. As judged by the presence of fixed examples (endogenous retroviruses), all jawed vertebrates live under threat of infection. In response, the host has developed mechanisms to prevent viral infections (1, 2). Forming part of the arsenal in the conflict with viruses are restriction factors, which inhibit various stages of the virus life cycle and act in a cell autonomous manner. Some of these, like TRIM5*α* (3), APOBEC3G (4) and SAMHD1 (5, 6), act at or before reverse transcription, while others, such as tetherin (7) and SERINC5 (8, 9), inhibit viral budding or fusion. In turn, viruses have developed measures to circumvent these blocks. The HIV-1 accessory genes *vif* and *vpu*, for example, specifically target APOBEC3G and tetherin for degradation, respectively (10, 11). Alternatively, sequence changes in the targets for restriction may allow virus escape.

The prototypic restriction factor, Fv1 (Friend virus susceptibility gene 1), was first described to protect laboratory mice against lethal infection by murine leukemia virus (MLV) (12, 13). Two alleles of *Fv1*, *Fv1^n^* and *Fv1^b^,* were originally described that act in a co-dominant fashion in heterozygous animals (14–16). We have since found that certain Fv1s from wild mice can additionally restrict non-MLV retroviruses (17). For example, Fv1 from *M. caroli* can restrict feline foamy virus (FFV), a spumavirus, and those from *M. spretus* and *M. macedonicus* were shown to restrict equine infectious anemia virus (EIAV), a lentivirus.

Fv1 targets the capsid (CA) protein present in the cytoplasm at a stage in retrovirus replication that is post-entry but before nuclear entry (18–21), binding to CA in the context of the hexametric lattice forming the viral core (22) and interfering with events downstream of reverse transcription (21). The specificity determinants of Fv1 map to the C-terminal part of the protein (CTD), indicating that this is the region that interacts with the viral capsid (23). The N-terminal region (NTD) of Fv1 contains a coiled coil motif that is involved in factor multimerization (22). This apparent means of binding has obvious parallels (24) to Trim5α, another CA-binding restriction factor, which forms a super-lattice over the viral core of infecting HIV-1 particles (25, 26).

Viruses breaching both adaptive and innate host defenses have the ability to significantly reduce the fitness of a host; viral burdens are, therefore, likely to have exerted substantial evolutionary pressures (27). Surveys of the variation of host genes influencing susceptibility to viruses provide useful information about the nature of the evolutionary race between viruses and their hosts and can illuminate mechanisms of viral escape. Positive selection of *Trim5α* has occurred for at least 30 million years (my) and has been shaped by the presence of lentiviruses (28–30). Similarly, we and others have uncovered equivalent forces acting upon *Fv1*, (17, 31–33) revealing a need for continuous or frequently reoccurring waves of retroviral infection in its maintenance over a ∼45 my lifetime (33).

To gauge whether retroviruses provide on-going selective pressure on the *Fv1* gene we have now set out to examine its variability within populations of wild mouse species from South East Asia (*M. caroli*, *M. cervicolor* and *M. cookii*). This work has revealed a duplication of the *Fv1* gene within this group of species, generating a novel gene we call *Fv7*. Both genes retain their expression capacity, show extensive variation and restriction assays reveal alleles with activity against spuma-, lenti-, and gammaretroviruses. The results of these studies suggest that restriction factor duplication can, at least in the case of *M. caroli*, allow a broadening of intrinsic immunity to confer simultaneous protection against multiple kinds of retrovirus.

## Results

### Duplication of *Fv1* in South East Asian mice

We have previously reported two Fv1 variants from *M. caroli*, differing in length by 8 amino acids (17). The longer variant (previously termed CAR1) restricted FFV and, to a lesser extent, prototypic foamy virus (PFV), while the shorter variant (CAR2), did not restrict any of the viruses in our panel. Both variants were cloned from CAROLI/EiJ mice purchased from the Jackson Laboratory. This strain has been maintained by closed colony breeding since 1994 and, as the mice were unlikely to be heterozygous, this led us to wonder if there could be two copies of the *Fv1* gene in *M. caroli*. This notion was encouraged by a separate report documenting two bands in a Southern hybridization experiment in which genomic DNA from *M. caroli* was probed with sequences corresponding to the 5’ end of Fv1 (31).

To investigate this possibility, we first made use of archived whole genome sequencing data made available under the Wellcome Sanger Institute’s Mouse Genomes Project, which includes CAROLI/EiJ (34, 35). Alignment of reads from the CAROLI/EiJ dataset to the C57BL/6J reference genome (GRCm38) revealed a doubling in the number of reads corresponding to *Fv1* compared to a C57BL/6NJ control sequenced as part of the same project (Fig 1A). This increase stretched both 5’ and 3’ of the *Fv1* locus and its range was used to conduct a focused breakpoint analysis. In addition to the canonical locus present on Chr 4, split-read and broken-pair data provided evidence of a second locus on Chr6 of CAROLI/EiJ and were used to design a typing PCR for the novel integration. Primers Chr6F and Chr6Rev anneal to the regions on Chr 6 flanking the novel insertion and, in the absence of an insertion, would yield a 900 bp PCR product. If the insertion, which is more than 3 kb in length, is present, there would be no product formed when employing a short extension time. PCR was performed using genomic DNAs from *M. caroli* and *M. spretus*, sourced from the Jackson Laboratory, as well as with those from wild-caught *M. fragilicauda*, *M. cookii*, *M. cervicolor* and *M. caroli* samples trapped in Thailand. Fragments of the predicted size for a ‘wild-type’ chromosomal region were observed for the reactions with Chr6F/Chr6Rev using DNAs from *M. spretus* and *M. fragilicauda*, confirming the absence of an insertion within these species (Fig 1B, top). Conversely, no PCR product was observed using this set of primers for *M. caroli*, *M. cookii* or *M. cervicolor*, consistent with an insertion between the sequences where the primers anneal on Chr 6 of these mice. A second primer pair, Chr4F/Chr6Rev, with one primer (Chr4F) annealing to a duplicated region of the *Fv1* locus was designed so that a fragment of 500 bp would be produced in the presence of an insert. Using this primer pair, PCR products were observed with DNAs from *M. caroli*, *M. cookii* and *M. cervicolor*, but not with *M. spretus* and *M. fragilicauda* (Fig 1B, bottom). The *M. caroli* samples yielded products around 200 bp larger than those of *M. cookii* and *M. cervicolor*, which, upon sequencing, was found to be due to the presence of a B1 short interspersed nuclear element (SINE) insertion upstream of the gene body.

**Fig 1.**
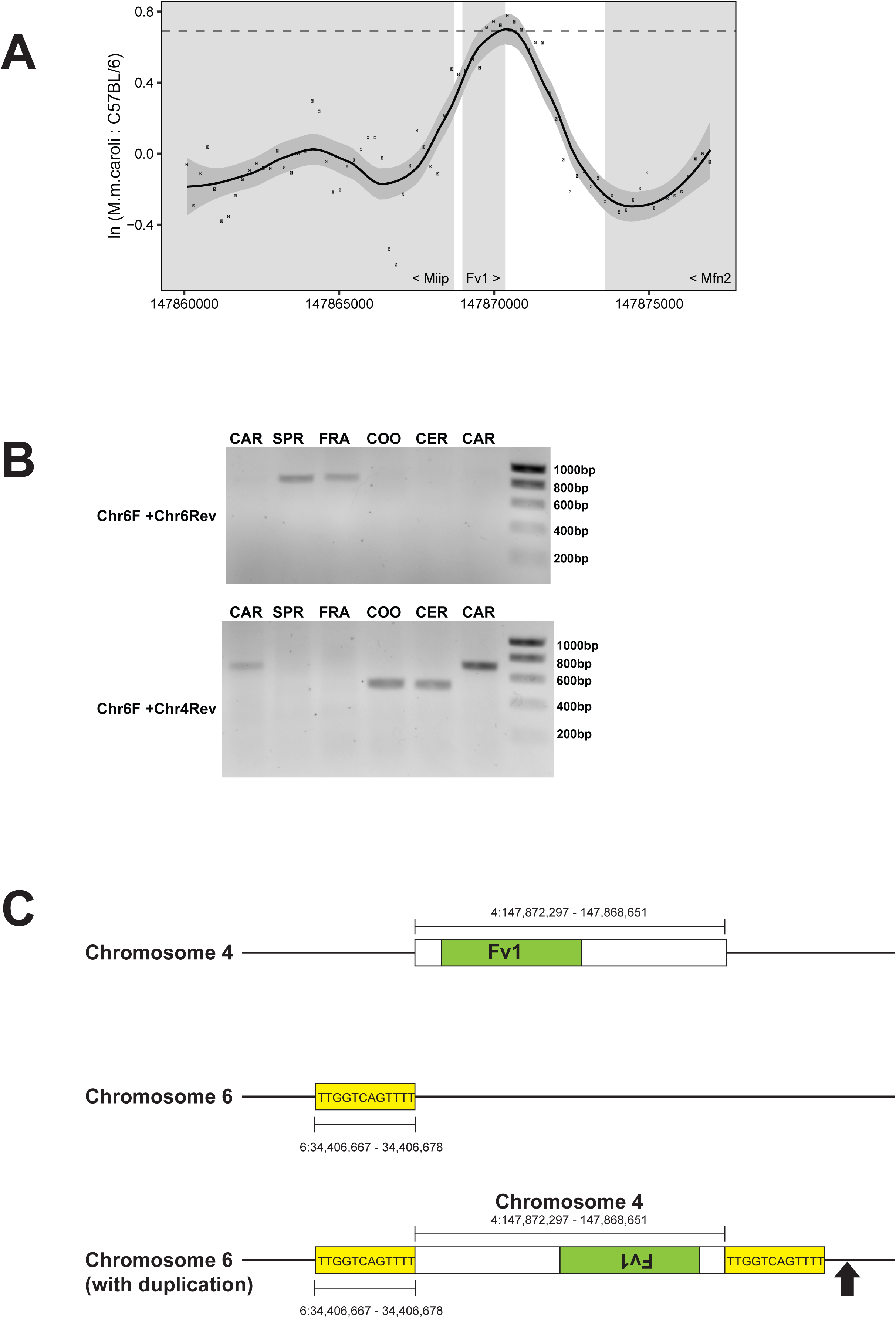
Duplication of *Fv1* in mice from South East Asia. **A**. Plot of the natural log of the ratio of the number of reads from the genomic sequence data of *M. caroli* to those of C57BL/6NJ mice along GRCm38 4:147,860,000-147,878,000. The shaded areas show the locations of the genes *Miip*, *Fv1* and *Mfn2* and the dashed line represents a ratio of 2. **B**. PCR strategy to confirm the insertion of an *Fv1* CDS on Chr 6 showing products of the primers detailed on the left. From left to right, CAR (CAROLI/EiJ from the Jackson Laboratory), SPR (SPRET/EiJ from the Jackson Laboratory), FRA (*M. fragilicauda* R7254), COO (*M. cookii* R7121), CER (*M. cervicolor* R6223), CAR (*M. caroli* R6321). **C**. A schematic representation of the insertion and the region of Chr duplicated. The direct repeat of the target sequence is highlighted in yellow and an arrow indicates the position of the *M. caroli* specific B1 insertion.

These results showed that a region of Chr 4 containing *Fv1* had been duplicated and inserted into Chr 6 of *M. caroli*, *M. cookii* and *M. cervicolor*, three closely related species of mice found in South East Asia. Together, these data provide compelling evidence that the gene duplication predated the divergence of these species and had not occurred during recent inbreeding.

Amplification and sequencing of the Chr 6 integration revealed the duplicated region corresponded to GRCm38 4:147868651–147872297 (3647 nts, extending 329 nts 5’ of the *Fv1* CDS and 1939 bp 3’ of the stop codon) (Fig 1C). The insertion was flanked by a 12 nt tandem site duplication (TSD), suggesting that the duplication occurred through long interspersed nuclear element (LINE)-mediated retrotransposition of an *Fv1* mRNA. Supporting this possibility, the duplicated region 3’ of the *Fv1* CDS and immediately preceding the TSD was terminated by a region of low complexity that did not share homology with the corresponding area of Chr 4. This region was dominated by p(A) stretches, likely evidence of the polyadenylation of the *Fv1* mRNA reverse transcribed by the LINE machinery.

The new locus, being separate from *Fv1*, was termed *Fv7*. The two previously studied variants from *M. caroli*, CAR1 and CAR2, could be assigned to *Fv1* and *Fv7*, respectively. By contrast, the *Fv1* gene we had previously isolated from *M. cervicolor* (CER) (17) was most probably a PCR-derived recombinant between the two genes.

### Genetic variation of *Fv1* and *Fv7* in the wild mouse populations of South East Asia

To test whether this gene duplication might play a role in protection against viruses endemic in South East Asia by allowing development of resistance to additional retroviruses, we set out to determine (a) the extent of sequence change in the novel gene, (b) whether it is transcribed, (c) whether sequence changes result in alterations of restriction specificity and (d) whether its presence allows a widening of protection to an extent not possible with *Fv1* alone.

To investigate the extent of natural variation in these genes, *Fv1* and *Fv7* from 44 mice (27 *M. cervicolor*, 7 *M. caroli*, 7 *M. cookii* and 3 *M. fragilicauda*), trapped in a range of locations across Thailand, were PCR-amplified and cloned using primers specific to the individual loci. Eight clones from each amplification were then sequenced. We identified 7 new *Fv1* and 9 *Fv7* variants from the *M. caroli* samples, 23 *Fv1* and 34 *Fv7* variants within *M. cervicolor* and 7 *Fv1* and 8 *Fv7* variants within *M. cookii*. There were 4 *Fv1* variants among the 3 *M. fragilicauda* samples. Details of the clones isolated are summarized in Table 1 and presented in Supporting Information (Appendix S1). Extensive sequence differences were visible, including point mutations, short insertions and deletions and a variety of duplications (Fig S1). This was consistent with the proposition that selection by diverse endemic viruses might be ongoing. It is noteworthy that *Fv1* and *Fv7* from *M. caroli* are more closely related to one another than to the *Fv1*s and *Fv7*s of *M. cookii* and *M. cervicolor* (Fig 2), perhaps suggesting that any selective forces are acting in a species-specific manner.

**Fig 2.**
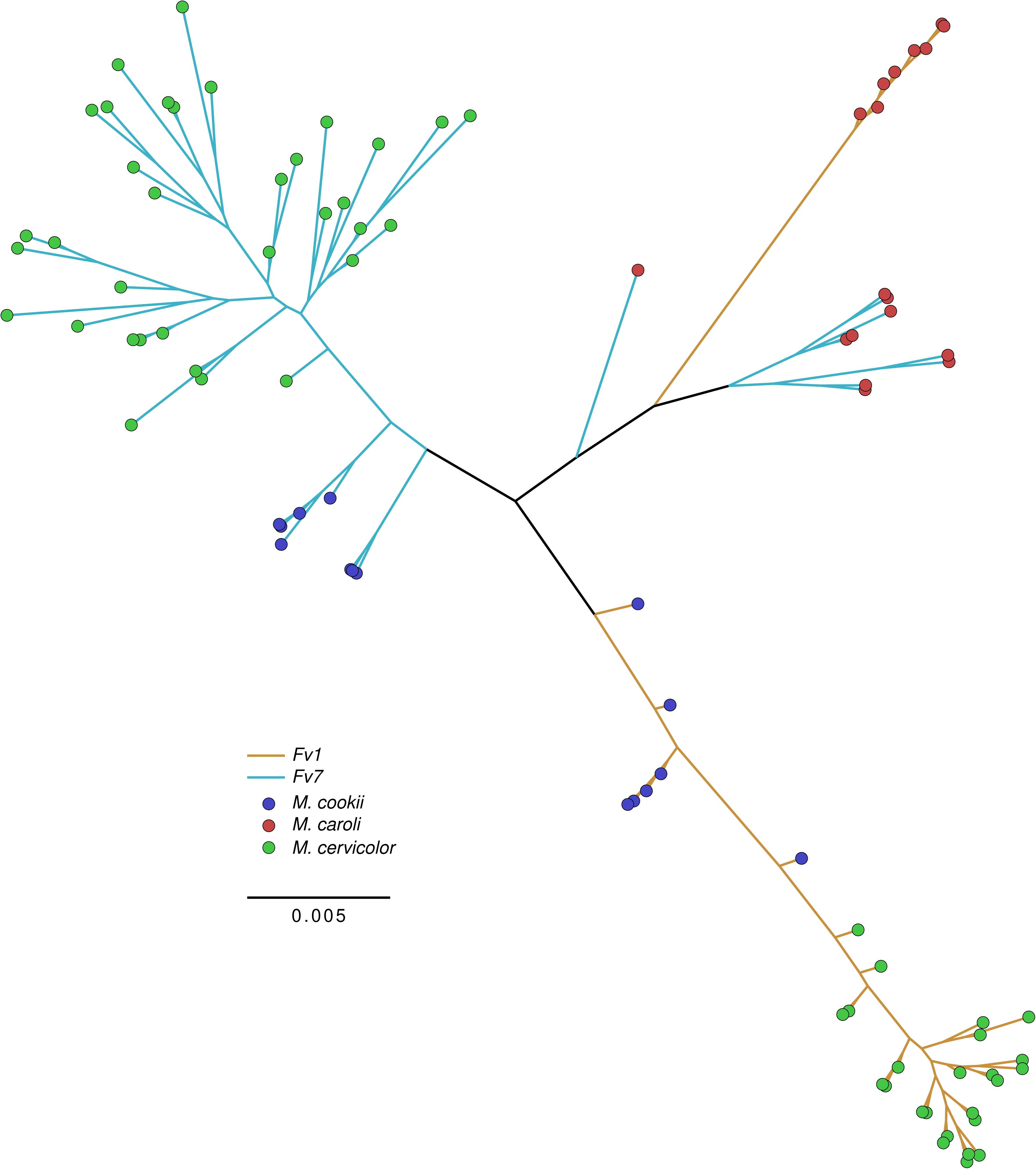
A phylogenetic tree of the *Fv1* and *Fv7* sequences. Unrooted maximum-likelihood tree of novel sequences prepared from an alignment with the variable tail removed (LogL -4107 under GTR+CAT, scale as substitutions per site).

**Table 1.** Fv1 and Fv7 variants present in mice from Thailand.

| ID | Species | Source / Location | Tissue | Variant |  |
| --- | --- | --- | --- | --- | --- |
|  |  |  |  | Fv1 | Fv7 |
| CAROLI/EiJ | *M. caroli* | Jackson Laboratory | Tail | 1 | 1 |
| R6321 | *M. caroli* | Kalasin | Spleen | 2, 3 | 2 |
| R6657 | *M. caroli* | Prachuapkirikhan | Spleen | 4 | 3, 4 |
| R6685 | *M. caroli* | Prachuapkirikhan | Spleen | 4, 5 | 3, 5 |
| R7195 | *M. caroli* | Nan | Liver | 6, 7 | 6 |
| R7225 | *M. caroli* | Nan | Liver | 6 | 7, 8 |
| R7236 | *M. caroli* | Nan | Liver | 6, 8 | 9, 7 |
| R7264 | *M. caroli* | Nakhon Ratchasima | Liver | 5 | 10 |
| R6223 | *M. cervicolor* | Kalasin | Spleen | 1, 2 | 1, (2) |
| R6243 | *M. cervicolor* | Kalasin | Spleen | 3*, 4 | 3, 4 |
| R6244 | *M. cervicolor* | Kalasin | Spleen | 5 | 3, 5 |
| R6254 | *M. cervicolor* | Kalasin | Spleen | 2 | 6, 7 |
| R6255 | *M. cervicolor* | Kalasin | Spleen | 3*, 6 | (8) |
| R6257 | *M. cervicolor* | Kalasin | Spleen | 1, 7 | 9, 10 |
| R6262 | *M. cervicolor* | Kalasin | Spleen | 2, 8 | 3, 11 |
| R6264 | *M. cervicolor* | Kalasin | Spleen | 4, 9 | 12, 13 |
| R6278 | *M. cervicolor* | Kalasin | Spleen | 3*, 10* | 14, 15 |
| R6279 | *M. cervicolor* | Kalasin | Spleen | 3* | 14, 15 |
| R6280 + | *M. cervicolor* | Kalasin | Spleen | 2 | 16, 17 |
| R6281 | *M. cervicolor* | Kalasin | Spleen | 11, 12 | (18), 19 |
| R6288 + | *M. cervicolor* | Kalasin | Spleen | 3*, 13 | 20 |
| R6293 | *M. cervicolor* | Kalasin | Spleen | 2 | 13, 21** |
| R6294 | *M. cervicolor* | Kalasin | Spleen | 14, 15 | 22, 23 |
| R6295 | *M. cervicolor* | Kalasin | Spleen | 4 | 17, 24 |
| R6297 + | *M. cervicolor* | Kalasin | Spleen | 1, 16 | 4, 21** |
| R6298 | *M. cervicolor* | Kalasin | Spleen | 13, 17 | 23 |
| R6320 | *M. cervicolor* | Kalasin | Spleen | 1, 16 | 10, 25 |
| R6682 | *M. cervicolor* | Prachuapkirikhan | Spleen | 18 | 26<sup>s</sup> |
| R6683 | *M. cervicolor* | Prachuapkirikhan | Spleen | 19 | 26<sup>s</sup>, 27 |
| R6684 + | *M. cervicolor* | Prachuapkirikhan | Spleen | 2 | 26<sup>s</sup> |
| R6701 | *M. cervicolor* | Prachuapkirikhan | Spleen | 2, 20* | 26<sup>s</sup>, 27 |
| R7255 + | *M. cervicolor* | Nakhon Ratchasima | Liver | 2, 21<sup>s</sup> | 28<sup>s</sup>, 29 |
| R7259 | *M. cervicolor* | Nakhon Ratchasima | Liver | 1, 2 | 30, 31 |
| R7262 + | *M. cervicolor* | Nakhon Ratchasima | Liver | 2, 22 | 14, 32 |
| R7263 | *M. cervicolor* | Nakhon Ratchasima | Liver | 23 | 33, 34 |
| R7121 | *M. cookii* | Tak | Liver | 1 | 1, (2) |
| R7180 | *M. cookii* | Nan | Liver | 2, (3) | (3), (2) |
| R7210 | *M. cookii* | Nan | Liver | 4 | (4) |
| R7237 | *M. cookii* | Nan | Liver | 4, (5) | (4) |
| R7238 | *M. cookii* | Nan | Liver | 4 | (4), (5) |
| R7239 | *M. cookii* | Nan | Liver | (3), (6) | 8, (6) |
| R7243 | *M. cookii* | Nan | Liver | 7 | (7) |
| R7254 | *M. fragicauda* | Nakhon Ratchasima | Liver | 1, 2 |  |
| R7260 | *M. fragicauda* | Nakhon Ratchasima | Liver | 3, 4 |  |
| R7261 | *M. fragicauda* | Nakhon Ratchasima | Liver | 3, 4 |  |

Variants in parentheses denote the presence of a premature stop codon that results in a truncated reading frame. <sup>s</sup> indicates an extended C-terminus, \* indicates internal repeats, \*\* indicates variants preceded by 2 residues before the consensus start codon, + indicates samples analyzed twice.

To confirm that the observed levels of variation were not an artefact of PCR amplification, we repeated the PCR, cloning and sequencing for 6 samples (Table 1), specifically including those with sequence duplications. In all cases, the clones sequenced exactly matched those seen in the initial analysis. Moreover, we only once observed more than two sequences per animal with a given primer pair; this one exception could be explained by recombination between *Fv1* and *Fv7* and was, therefore, excluded from all further analysis and is not reported here. Thus, the variation seen truly reflects genetic variation in the natural population, with heterozygosity frequently observed for both genes.

A number of samples encoded frameshifted or truncated proteins. These were particularly common among the *M. cookii* samples; indeed, when amplified with *Fv7*-specific primers, only Fv7COO1 and 8 encoded an open reading frame (Table 1). Considering allelic variation, all samples revealed at least one variant with a five-nucleotide deletion resulting in a frameshift and a truncated protein of 404 amino acids. Similarly, three of the same samples encoded at least one truncated Fv1 protein (Table 1). We have previously shown that truncation of *Fv1* to 410 amino acids results in complete loss of restriction activity (23), making functionality of the *Fv7* and *Fv1* truncations improbable. Nevertheless, all *M. cookii* examined contained at least one allele of either gene with intact coding potential.

All *Fv1* genes from *M. cervicolor* and *M. cookii* mice contained a B1 SINE inserted at an identical position within the 3’ region of the open reading frame, resulting in a protein with a shortened C-terminus (Fig S2). Consistent with current estimations of *Mus* phylogeny (36, 37), the absence of the B1 insert in *M. caroli* indicated that the transposition event occurred after the separation of *M. caroli* and the common ancestor of *M. cervicolor* and *M. cookii*. The *Fv1* gene of *M. fragilicauda* also contained a B1 SINE apparently at the same position, yet other sequence differences, as well as the earlier divergence of this species from the M. *caroli/cervicolor/cookii* clade, indicate the independent acquisition of this B1 insertion. We have previously reported the presence of independent B1 insertions in 3 other species of mice (*M. famulus*, *M. minutoides* and *M. platythrix*) within a few nucleotides of those seen here (17). Thus, this provides further suggestive evidence that minimizing the length of the C-terminus may provide enhanced restriction properties and outlines a strong, convergent, role for exploitation of the mobility of B1 SINEs in realizing this adaptation across species.

Overall, our results showed the presence of a rich diversity of *Fv1* and Fv7 sequences in the wild mouse populations of South East Asia. Accumulation and fixation of single nucleotide polymorphisms and co-option of retroelement mobility as a means of separation and diversification of the protein sequences of the two genes suggests a strong role for positive selection in their evolution.

### Expression of Fv1 and Fv7

Despite its retroviral origin, a viral long terminal repeat on the 5’ side of *Fv1* is not present within *Mus*, although degraded fragments can be noted in more distantly-related genera (33). In its absence, transcription was, therefore, thought to be driven from bidirectional promoter activity of the adjacent antisense gene, *Miip*. Neither the exact promoter region nor the point of transcript initiation has been defined, however, raising the question of whether the duplicated region on Chr 6 retained the potential to drive expression of *Fv7*. Hence, we set out to map the promoter region of the parental *Fv1* locus.

Fragments containing increasing lengths of the region 5’ of the *Fv1* CDS on Chr 4 were cloned into pGL4.10, ahead of a promoterless *Luc* gene (Fig 3A). The constructs were transfected into *M. dunni* tail fibroblast (MDTF) cells and the luciferase activities measured. Relative luciferase activity first increased above the background of the promoterless plasmid with the construct containing 250 nts upstream of the *Fv1* start codon and further increased with inclusion of regions up to 300 and 350 nts (Fig 3B). Activities observed were, nevertheless, less than 25% of those obtained with the SV40 promoter of the pGL4.13 control, consistent with the low levels of endogenous *Fv1* expression previously described (38).

**Fig 3.**
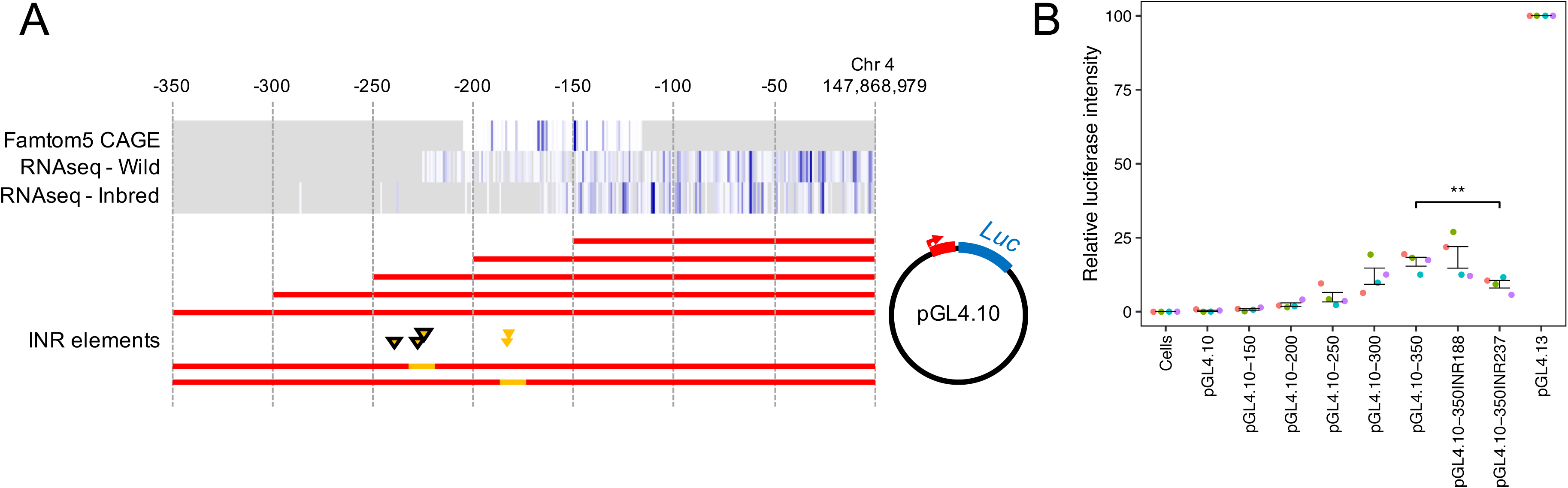
Analysis of *Fv1* transcription. **A**. Schematic of plasmid constructs produced to test for promoter activity within the region 5’ of the *Fv1* CDS. Heatmaps (white to blue, grey indicates no data) detail the position of identified transcription start sites within FANTOM5 CAGE data and, separately, for the positions of Illumina RNAseq reads from wild mice and inbred strains. Yellow triangles indicate the positions of INR element predictions (black borders indicate high confidence predictions) and accompanying yellow highlights indicate regions mutated to adenine. **B**. Relative luciferase intensity for control (Cells, pGL4.10, and pGL4.13) and experimental plasmids. Data are 4 independent experiments analyzed in triplicate, and normalized to the intensity of pGL4.13. A paired two-tailed student’s T-test was used to determine significant reduction for 350INR237 (350INR188 was not altered).

To better determine potential points of transcriptional initiation, we extracted cap analysis gene expression (CAGE) data from the FANTOM5 project (39) for endogenous *Fv1* expression. Dispersed transcription start sites were identified between 80 to 170 nucleotides 5’ of the Fv1 ATG (Fig 3A) but did not represent the full extent of transcription within the region determined in a complementary analysis of RNAseq reads from 9 inbred laboratory mouse strains (accession ERP000614) (40), which identified dispersed points of initiation beyond 200 nts 5’ of the *Fv1* ATG (Fig 3A). Three partially-overlapping high-confidence initiator (INR) element predictions with 94- 97% satisfaction of an INR position weight matrix (PWM) model (41) could be determined that supported the area additionally identified within the RNAseq data (Fig 3A), whereas only two, overlapping, low-confidence (81% PWM model satisfaction), predictions could be made within the areas identified by CAGE (Fig 3A). Whilst initiation is certainly dispersed, therefore, we sought to investigate any specific contributions of these regions with mutated constructs. Mutation of the high-confidence INR at 237 nucleotides 5’ of the *Fv1* ATG produced a significant reduction in luciferase expression (Fig 3B), suggesting its likely involvement in *Fv1* transcription. By contrast, disruption of the low-confidence INR elements at 188 seemed unimportant.

The promoter region of *Fv1* is likely cryptic and transcriptional initiation can be seen to occur across of range of sites. Nevertheless, these data confirmed that a region sufficient for *in vitro* expression had been retrotransposed onto Chr 6. To confirm explicitly that co-expression of *Fv1* and *Fv7* could occur *in vivo*, we pooled and analyzed available RNAseq data for *M. caroli* (accessions ERP001409 and ERP005559) (35, 42). As similarity of the genes complicated expression assessment due to ambiguity in the assignation of multi-mapping reads, reads at positions where the two genes consistently varied (‘trunk’ differences) were manually inspected and assigned. This process revealed reads originating from both genes, confirming that the region retrotransposed to Chr 6 and present in *M. caroli*, *M. cervicolor* and *M. cookii*, is sufficient for *in vivo* expression and that co-expression occurs naturally.

### Restriction specificities of cloned Fv1s and Fv7s

As previously hypothesized, high levels of sequence variation within *Fv1* may be due to selection by a range of retroviruses, which are likely to have contributed to maintenance and diversification of the gene (33). The extensive variation among the Fv1 and Fv7 sequences observed here thus led us to wonder if they were capable of recognizing multiple viruses. We therefore tested a subset of these novel sequences for their ability to restrict a comprehensive panel of retroviruses (Tables 2 and 3).

**Table 2.** Restriction of MLVs by selected Fv1s and Fv7s.

| **Variant** | **N-MLV** | **B-MLV** | **Mo-MLV** | **NR-MLV** |
| --- | --- | --- | --- | --- |
| Fv1CAR2 | 1.16 ± 0.02 | 0.34 ± 0.02 | 1.01 ± 0.02 | 1.16 ± 0.10 |
| Fv1CAR3 | 1.10 ± 0.14 | 0.34 ± 0.02 | 0.83 ± 0.06 | 1.12 ± 0.12 |
| Fv1CAR4 | 1.19 ± 0.04 | 0.53 ± 0.08 | 1.08 ± 0.05 | 1.12 ± 0.12 |
| Fv1CER1 | 0.35 ± 0.02 | 0.03 ± 0.01 | 1.06 ± 0.12 | 0.43 ± 0.05 |
| Fv1CER2 | 0.04 ± 0.01 | 0.04 ± 0.01 | 1.03 ± 0.06 | 0.05 ± 0.01 |
| Fv1CER19 | 0.09 ± 0.03 | 0.06 ± 0.01 | 1.03 ± 0.03 | 0.09 ± 0.01 |
| Fv1CER21 | 0.13 ± 0.01 | 0.12 ± 0.01 | 1.06 ± 0.01 | N.D |
| Fv1CER22 | 1.16 ± 0.01 | 1.17 ± 0.12 | 1.16 ± 0.05 | 1.11 ± 0.08 |
| Fv1COO1 | 1.13 ± 0.07 | 0.15 ± 0.01 | 1.23 ± 0.10 | 1.19 ± 0.21 |
| Fv1COO2 | 0.07 ± 0.01 | 0.06 ± 0.01 | 0.33 ± 0.05 | 0.07 ± 0.01 |
| Fv1COO4 | 1.11 ± 0.04 | 0.03 ± 0.01 | 1.14 ± 0.03 | 1.21 ± 0.07 |
| Fv1COO7 | 0.03 ± 0.01 | 0.03 ± 0.01 | 0.13 ± 0.06 | 0.03 ± 0.01 |
| Fv1FRA1 | 0.09 ± 0.01 | 0.09 ± 0.01 | 0.08 ± 0.01 | 0.09 ± 0.00 |
| Fv1FRA2 | 0.09 ± 0.00 | 0.09 ± 0.01 | 0.10 ± 0.01 | 0.10 ± 0.01 |
| Fv7CAR2 | 1.12 ± 0.17 | 0.70 ± 0.16 | 0.87 ± 0.09 | 1.16 ± 0.02 |
| Fv7CAR3 | 1.18 ± 0.28 | 0.74 ± 0.28 | 0.82 ± 0.09 | 1.23 ± 0.20 |
| Fv7CAR4 | 0.70 ± 0.02 | 1.03 ± 0.24 | 0.86 ± 0.07 | 1.14 ± 0.13 |
| Fv7CER26 | 1.16 ± 0.04 | 1.14 ± 0.03 | 1.16 ± 0.02 | 1.09 ± 0.03 |
| Fv7CER27 | 0.41 ± 0.05 | 1.41 ± 0.17 | 1.18 ± 0.03 | 1.15 ± 0.03 |
| Fv7CER29 | 1.06 ± 0.01 | 1.14 ± 0.06 | 1.12 ± 0.03 | 1.14 ± 0.06 |
| Fv7CER30 | 0.33 ± 0.04 | 1.22 ± 0.08 | 1.16 ± 0.08 | 1.18 ± 0.01 |
| Fv7CER31 | 0.37 ± 0.05 | 1.23 ± 0.05 | 1.14 ± 0.02 | 1.16 ± 0.24 |
| Fv7COO1 | 0.02 ± 0.01 | 1.04 ± 0.35 | 1.17 ± 0.25 | 1.21 ± 0.23 |
| Fv7COO8 | 0.06 ± 0.04 | 0.97 ± 0.22 | 0.88 ± 0.34 | 1.13 ± 0.10 |

Restriction assays were performed as described in the Materials and Methods. Values are the means and standard deviations of at least 4 experiments.

**Table 3.**
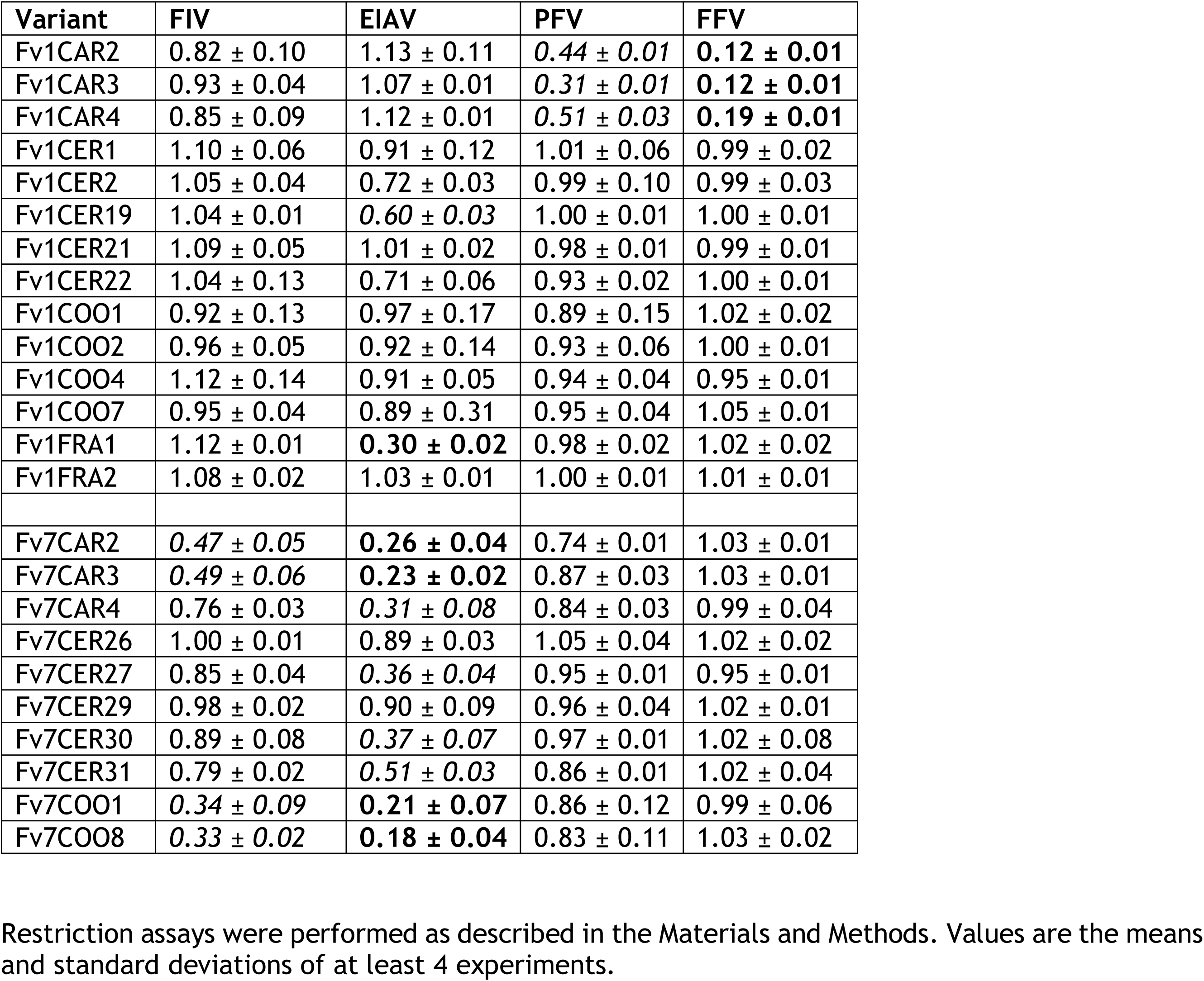
Restriction of non-MLV retroviruses by Fv1s and Fv7s.

| **Variant** | **FIV** | **EIAV** | **PFV** | **FFV** |
| --- | --- | --- | --- | --- |
| Fv1CAR2 | 0.82 ± 0.10 | 1.13 ± 0.11 | 0.44 ± 0.01 | **0.12 ± 0.01** |
| Fv1CAR3 | 0.93 ± 0.04 | 1.07 ± 0.01 | 0.31 ± 0.01 | **0.12 ± 0.01** |
| Fv1CAR4 | 0.85 ± 0.09 | 1.12 ± 0.01 | 0.51 ± 0.03 | **0.19 ± 0.01** |
| Fv1CER1 | 1.10 ± 0.06 | 0.91 ± 0.12 | 1.01 ± 0.06 | 0.99 ± 0.02 |
| Fv1CER2 | 1.05 ± 0.04 | 0.72 ± 0.03 | 0.99 ± 0.10 | 0.99 ± 0.03 |
| Fv1CER19 | 1.04 ± 0.01 | 0.60 ± 0.03 | 1.00 ± 0.01 | 1.00 ± 0.01 |
| Fv1CER21 | 1.09 ± 0.05 | 1.01 ± 0.02 | 0.98 ± 0.01 | 0.99 ± 0.01 |
| Fv1CER22 | 1.04 ± 0.13 | 0.71 ± 0.06 | 0.93 ± 0.02 | 1.00 ± 0.01 |
| Fv1COO1 | 0.92 ± 0.13 | 0.97 ± 0.17 | 0.89 ± 0.15 | 1.02 ± 0.02 |
| Fv1COO2 | 0.96 ± 0.05 | 0.92 ± 0.14 | 0.93 ± 0.06 | 1.00 ± 0.01 |
| Fv1COO4 | 1.12 ± 0.14 | 0.91 ± 0.05 | 0.94 ± 0.04 | 0.95 ± 0.01 |
| Fv1COO7 | 0.95 ± 0.04 | 0.89 ± 0.31 | 0.95 ± 0.04 | 1.05 ± 0.01 |
| Fv1FRA1 | 1.12 ± 0.01 | **0.30 ± 0.02** | 0.98 ± 0.02 | 1.02 ± 0.02 |
| Fv1FRA2 | 1.08 ± 0.02 | 1.03 ± 0.01 | 1.00 ± 0.01 | 1.01 ± 0.01 |
| Fv7CAR2 | 0.47 ± 0.05 | **0.26 ± 0.04** | 0.74 ± 0.01 | 1.03 ± 0.01 |
| Fv7CAR3 | 0.49 ± 0.06 | **0.23 ± 0.02** | 0.87 ± 0.03 | 1.03 ± 0.01 |
| Fv7CAR4 | 0.76 ± 0.03 | 0.31 ± 0.08 | 0.84 ± 0.03 | 0.99 ± 0.04 |
| Fv7CER26 | 1.00 ± 0.01 | 0.89 ± 0.03 | 1.05 ± 0.04 | 1.02 ± 0.02 |
| Fv7CER27 | 0.85 ± 0.04 | 0.36 ± 0.04 | 0.95 ± 0.01 | 0.95 ± 0.01 |
| Fv7CER29 | 0.98 ± 0.02 | 0.90 ± 0.09 | 0.96 ± 0.04 | 1.02 ± 0.01 |
| Fv7CER30 | 0.89 ± 0.08 | 0.37 ± 0.07 | 0.97 ± 0.01 | 1.02 ± 0.08 |
| Fv7CER31 | 0.79 ± 0.02 | 0.51 ± 0.03 | 0.86 ± 0.01 | 1.02 ± 0.04 |
| Fv7COO1 | 0.34 ± 0.09 | **0.21 ± 0.07** | 0.86 ± 0.12 | 0.99 ± 0.06 |
| Fv7COO8 | 0.33 ± 0.02 | **0.18 ± 0.04** | 0.83 ± 0.11 | 1.03 ± 0.02 |

Restriction assays were performed as described in the Materials and Methods. Values are the means and standard deviations of at least 4 experiments.

Contrary to our previous report analyzing Fv1CAR1 (then termed CAR1) (17), which determined no activity against MLV, all three Fv1CARs tested here gave partial restriction of B-MLV (Table 2). Comparison of Fv1CAR1 and Fv1CAR2 showed three amino acid differences and exchange of a single residue, Fv1CAR428, restored activity against B-MLV without affecting that seen against FFV (Table 3, Fig S3). The remaining Fv1s restricted a wider array of MLVs (Table 2). Fv1CER1 and Fv1CER2 as well as Fv1COO4 and Fv1COO7 differed in their ability to recognize NR-MLV; in both cases this difference could be mapped to single amino acids (Fv1CER428 and Fv1COO268) (Fig S4). It is noteworthy that Fv1CER recognized NB-MLV whereas Fv1COO did not. On several occasions, e.g. Fv1CAR2, 3, 4 and Fv1COO1, reduced restriction activity seemed correlated with a longer CTD and it would be interesting to test the effect of artificially truncating the Fv1s from Fv1CAR2 and Fv1COO1 in a manner analogous to that seen with the B1 repeat in Fv1CER. Thus, most Fv1s showed at least limited effects on MLV.

Consistent with our previous study (17), Fv1CAR2, Fv1CAR3 and Fv1CAR4 all restricted FFV fully and PFV to a lesser extent (Table 3). None of the other factors tested had this effect (Table 3). All the Fv1s from *M. caroli* in this study contained the determinants (K348, Y351) previously shown (17) to be responsible for foamy virus restriction, implying that all variants will likely restrict FFV and PFV. This might suggest the widespread exposure to FFV-like viruses in the current *M. caroli* population in Thailand – individual samples coming from Prachuapkirikhan in the south, Kalasin in the east and Nan in the north of Thailand (Table 1).

Interestingly, at least one of the Fv1s from *M. fragilicauda* and several of the Fv7s from *M. caroli*, *M. cervicolor* and *M. cookii* exhibited EIAV restriction (Table 3). We have previously mapped the ability of *M. spretus Fv1* to recognize EIAV to a R268C change (17) and, similarly here, we find that a C is again present at the analogous position in Fv1FRA1, which restricts, but not in Fv1FRA2, which does not. This amino acid is not found in any of the restricting Fv7s, however, where the changes responsible for restriction remain elusive. These active Fv7s further differ from Fv1FRA1 in their ability to partially restrict FIV (Table 3). We had previously cloned *Fv7* from *M. caroli* (then termed CAR2) (17), which differs from the seven Fv7s cloned here by the presence of a unique G at residue 351 in place of an E. Introduction of E351G restores the ability to restrict EIAV (Fig S5). These results would be consistent with the presence of an FIV- or EIAV-like virus endemic in the area and selecting for the observed restriction activity.

### Combining EIAV and FFV restriction

Two individual *M. caroli* samples from different locations, identifiers R6321 and R6657 (Table 1), both carried an Fv1 that restricted FFV and MLV and an Fv7 that restricted EIAV and FIV. Indeed, based on the sequences described here, it is likely that this applies to all *M. caroli* sampled, suggesting conference of a certain selective advantage. This raised the question as to whether such differing restriction profiles could be achieved within a single gene or whether gene duplication and diversification was required to achieve such broad recognition.

To examine this idea further, we first tried creating a single restriction factor with the ability to recognize both lenti- and foamy viruses. Introduction of the residues conferring FFV restriction (17) into the Fv7s recognizing EIAV achieved only a very weak restriction in Fv7CAR2 and Fv7CAR3 but not in Fv7CER27 and Fv7COO8 (Table 4). Further, in all cases, the EIAV restriction was abolished. The alternate introduction of FFV determinants into Fv1^n^ carrying R268C again proved unsuccessful (Table 4).

**Table 4.** Incompatibility of FFV and EIAV restriction determinants.

| **Variant** | **FFV** | **EIAV** | **N-MLV** | **B-MLV** |
| --- | --- | --- | --- | --- |
| CAR1 | 0.15 ± 0.06 | 1.10 ± 0.14 | 1.09 ± 0.09 | 1.08 ± 0.06 |
| FV7CAR2 | 1.01 ± 0.05 | 0.24 ± 0.07 | 1.04 ± 0.07 | 1.08 ± 0.05 |
| Fv7CAR2 E349K S352Y | 0.68 ± 0.18 | 0.99 ± 0.08 | 1.12 ± 0.03 | 1.05 ± 0.04 |
| Fv7CAR3 | 1.00 ± 0.04 | 0.20 ± 0.02 | 1.03 ± 0.10 | 1.09 ± 0.06 |
| FvCAR3 E349K S352Y | 0.59 ± 0.11 | 0.93 ± 0.11 | 1.06 ± 0.03 | 1.04 ± 0.09 |
| Fv7CER27 | 1.01 ± 0.06 | 0.36 ± 0.05 | 0.46 ± 0.04 | 1.12 ± 0.05 |
| Fv7CER27 E349K S352Y | 1.04 ± 0.03 | 1.06 ± 0.03 | 1.07 ± 0.05 | 0.94 ± 0.07 |
| Fv7COO8 | 0.99 ± 0.06 | 0.22 ± 0.04 | 0.11 ± 0.04 | 1.08 ± 0.09 |
| Fv7COO8 E349K S352Y | 1.02 ± 0.01 | 1.00 ± 0.12 | 1.08 ± 0.03 | 1.11 ± 0.05 |
| Fv1n | 1.04 ± 0.03 | 1.21 ± 0.05 | 1.12 ± 0.02 | 0.09 ± 0.02 |
| Fv1n R268C | 1.00 ± 0.02 | 0.23 ± 0.02 | 0.15 ± 0.03 | 0.16 ± 0.02 |
| Fv1n E349K S352Y | 0.13 ± 0.03 | 1.12 ± 0.08 | 1.07 ± 0.04 | 0.04 ± 0.01 |
| Fv1n R268C E349K S352Y | 0.99 ± 0.04 | 1.23 ± 0.04 | 0.45 ± 0.02 | 0.21 ± 0.06 |

Restriction assays were performed as described in the Materials and Methods. Values are the means and standard deviations of at least 4 experiments.

Alternatively, and considering that Fv1 activity was initially described as co-dominant, we sought to test whether co-expression of Fv1s with different restriction specificities could protect a cell against multiple viruses. For this purpose, a three-color flow cytometry restriction assay was established in which permissive MDTFs were transduced with two retroviral vectors expressing different restriction factors together with either EYFP or mScarlet, so that cells which were transduced with one restriction gene were either yellow or red while those containing both restriction genes were doubly labelled. The mixed population was then challenged with tester viruses carrying an EGFP construct. Thus, each population could be individually identified by FACS, allowing infection susceptibility to be scored as the percentage of green cells within each population. Restriction was expressed as the ratio of the percentage of infection in cells containing restriction factors (either yellow, red or yellow and red) to those which did not (unlabelled).

Fv1^n^ or Fv1^b^ alone provided strong restriction activity against B-MLV or N-MLV, respectively (Fig 4A), however their co-expression in the same cell led to a slight reduction in restriction of either virus. Nevertheless, significant restrictions of both N-MLV and B-MLV were still observed, supporting the determination of co-dominance for these alleles (16). By contrast, in cells expressing both Fv1CAR and Fv7CAR, complete loss of restriction of both FFV and EIAV was observed, indicating apparent interference between the co-present factors. Equivalent interference has previously been reported between the TRIM5α proteins of Human and rhesus macaque and, similarly, between Human TRIM5α and owl monkey TRIMCyp, a TRIM5-cyclophillin fusion (43).

**Fig 4.**
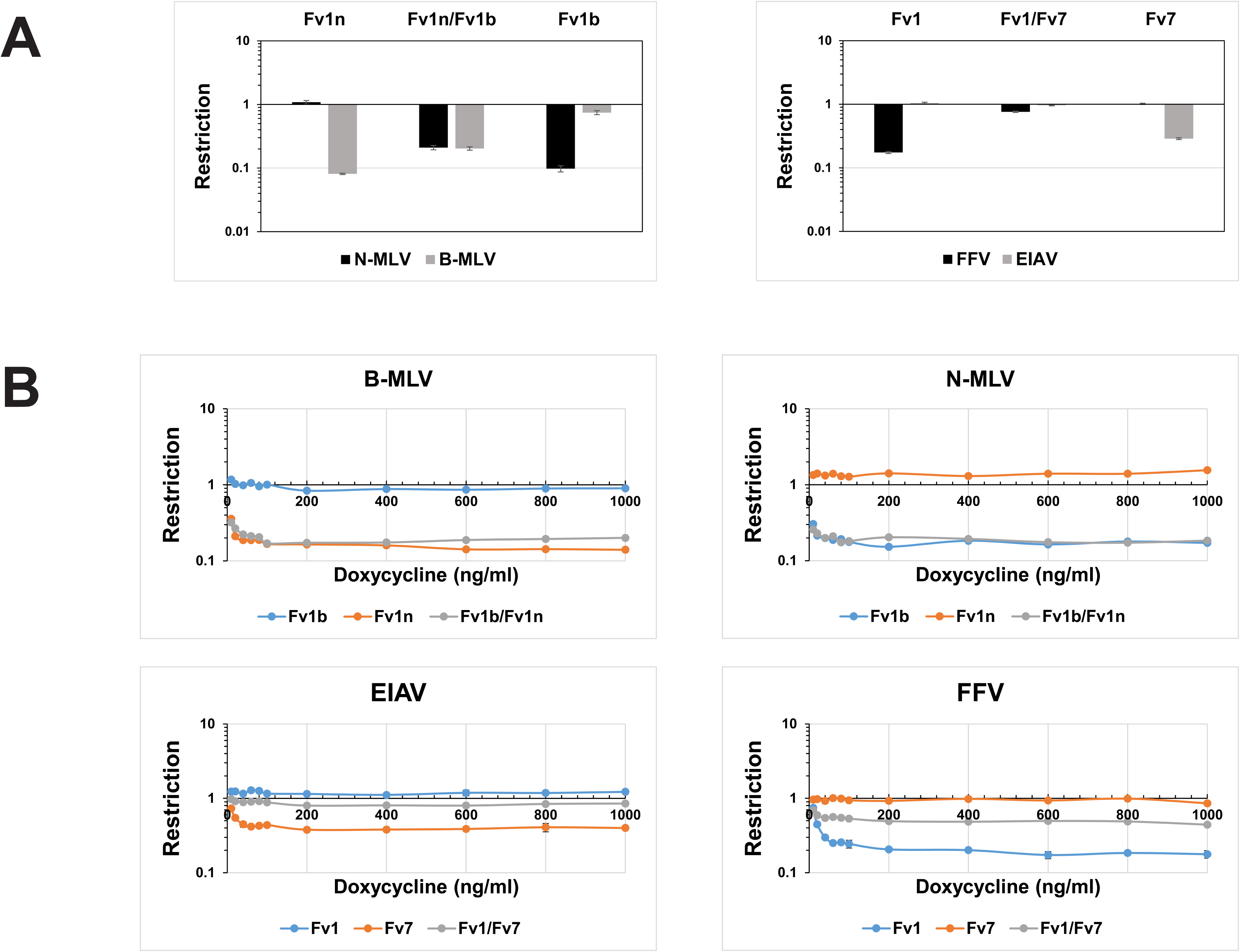
Co-expression of Fv1CAR and Fv7CAR. **A**. Expression from a retroviral promoter. MDTFs were transduced to co-express Fv1^b^/mScarlet and Fv1^n^/EYFP (left) or Fv1CAR/mScarlet and Fv7CAR/EYFP (right) and challenged with either N-MLV, B-MLV, FFV or EIAV carrying the EGFP gene. **B**. Expression from an inducible promoter over a range of induction. The same combinations of fluorescence genes and Fv1 or Fv7 were placed under the control of a doxycycline inducible promoter in retroviral vectors that have been previously described and used to transduce R18 cells. The cells were induced with doxycycline concentrations from 10 ng/ml to 1000 ng/ml for 24 hours before challenge. In both A and B, restriction is expressed as the ratio of the percentages of cells containing restriction factor(s) that were infected to those of cells that did not contain restriction factor and were infected.

We have previously noted that levels of *Fv1* expression can impact determination of restriction activities, however, as endogenous levels of *Fv1^n^* and *Fv1^b^* are very low (38). As the first set of experiments was performed with vectors expressing the restriction factors from retroviral promoters, it was possible, therefore, that the reduction in restriction activities observed was due to their relative overexpression. To test this hypothesis, we repeated the assay using inducible promoters to express the restriction genes. As before, the ability of Fv1^n^ or Fv1^b^ to restrict either B-MLV or N-MLV, respectively, was almost identical whether they were present individually or together over a wide range of doxycycline concentrations (Fig 4B). Across all levels of induction, however, co-expression of Fv1CAR and Fv7CAR abolished anti-EIAV activity and markedly reduced anti-FFV activity (Fig 4B). Even at physiological levels of expression, co-expression of these factors resulted in interference, therefore, suggesting that the ability of different restriction factors to inhibit diverse viruses in the same cell may be dependent on the nature of their viral targets.

## Discussion

Diverse retroviruses have undoubtedly exerted sustained selection pressures through both human (28–30) and murid (33) evolution. For both, a variety of ecological considerations – population density and exposure to other co-endemic species, for example – have influenced exposure to circulating retroviruses. These, as well as other spaciotemporal factors have likely contributed in the a wide array of restriction profiles now visible across species of *Mus* (17). Previous work (17), as well as experiments within the present study, indicate limitations in the ability of differing Fv1-based restriction profiles to be additively merged within single proteins, however. For example, attempts to generate an Fv1 that restricts both FFV and EIAV have not proved successful. Such limitations, possibly visualized as separate peaks within an evolutionary landscape, potentially limit overall restriction plasticity. We now detail the first example of *Fv1* duplication and the acquisition of differing restriction profiles within Fv7 and Fv1 as a means of enhancing restriction range.

Given the presence of a 12 nt tandem site duplication and the integration of a non-templated region likely resulting from mRNA polyadenylation (44), it is probable that the *Fv7* locus on Chr 6 results from LINE-mediated retrotransposition. However, the definitive hallmark of retrogenes, exon merger as a result of splicing (45), is missing because *Fv1* comprises a single exon. The region duplicated contains 329 nt of sequence upstream of the *Fv1* CDS on Chr 4, thereby encompassing sufficient sequence for promoter activity *in vitro*. Analysis of published RNAseq data from *M. caroli* confirms expression of both loci.

*Fv7* within *Mus caroli*, *M. cervicolor* and *M. cookii* encodes a protein with equivalent functional properties to Fv1. Indeed, where we have successfully mapped certain restriction activities to specific amino acids, all fall within the previously defined variable regions of Fv1 responsible for restriction of different viruses (17, 33). Examples include Fv1CAR1 residue 428 (Fig S3), Fv1COO4 residue 268 (Fig S4) and Fv7CAR1 residue 351 (Fig S5). However, it is noteworthy that residues 358 and 399 of *M. musculus* Fv1, key for distinguishing between N-MLV and B-MLV in Fv1^n^ and Fv1^b^ (23), are identical across all Fv1s and Fv7s cloned here, despite the differences in MLV restriction visible at both the individual and species level (Table 2).

Our attempts *in vitro* to introduce FFV- and EIAV-restricting Fv1s and Fv7s into the same cell, even at endogenous expression levels, have not resulted in dual restriction (Fig 4). Fv1 restriction is thought to involve formation of a multimeric lattice around incoming virions (24) in a manner analogous to the TRIM5*α* complexes engulfing incoming retroviruses (26, 46). The incoming cores of lentiviruses and foamy viruses have different arrays of Gag proteins and it is possible that, at least within our assay system, formation of mixed Fv1 complexes does not result in stable binding. Nevertheless, it is clear that the generation, genetic fixation and maintenance of different activities within these species has taken place – a process undeniably requiring positive selection by conference of resistance to viral infection. This represents a certain paradox, therefore. Differential expression of the two genes in different tissues might provide one explanation but further characterization of the transcriptional profiles of the two genes would be required to shed light on how this might occur. In support of such an explanation, it is noteworthy that, within *M. caroli,* the *Fv7* locus has accumulated an in-frame B1_Mus2 SINE element upstream of the CDS, one of only few families showing potential links to gene regulation (47). Whether the B1 element inserted here acts to alter Fv1 production remains to be determined.

Across the species surveyed, the Fv1 and Fv7 proteins show substantial sequence variation and adaptation to recognize viruses of different genera. Unfortunately, a sparsity of whole genome sequencing data from multiple individuals of diverse *Mus* species prevents the comparison of relative rates of polymorphism. Nevertheless, an indicative comparison to sequences previously determined for *M. domesticus* and *M. musculus* (17, 31, 48) suggests a higher extent of sequence variation than might be expected. Behind the levels of allelism detailed, it seems probable that manifold viruses circulate within Thai mice. Though the viruses driving these changes have not been identified, on the assumption that the driver viruses resemble those defining the observed activities, it would seem reasonable to conclude that Thai mice have been exposed to both foamy and lentiviruses. Further, these viruses must have been sufficiently pathogenic to drive the generation and fixation of novel resistance genes. To the best of our knowledge, no mouse-tropic foamy or lentiviruses have ever been described and, given the potential for murids to act as vector species (49), a search for such viruses might be of considerable interest. Finally, we would like to suggest that Fv1 is not alone in its evolutionary plasticity. Other antiviral factors, for example members of the Trim family (50, 51), protecting against other types of virus (52), seem likely to exhibit significant sequence variation driven by circulating viruses.

## Materials and Methods

### Mice

Wild mice were trapped in different provinces of Thailand as listed in Table 1. Approval notices for trapping and investigation of rodents were provided by the Ethical Committee of Mahidol University, Bangkok, Thailand, number 0517.1116/661; animals were treated in accordance with the guidelines of the American Society of Mammalogists, and within the European Union legislation guidelines (Directive 86/60/EEC). Spleens or livers were removed and frozen for later DNA extraction using the Qiagen DNeasy Blood and Tissue kit according to the manufacturer’s instructions. Species identification was confirmed by PCR with a mitochondrial DNA bar-coding method. Briefly, a segment of the cytochrome oxidase subunit 1(COI) gene was amplified gDNA using the primers BatL5310 (5’ CCTACTCRGCCATTTTACCTATG 3’) and R6036R (5’ ACTTCTGGGTGTCCAAAGAATCA 3’). The sequence of the PCR fragment was then used in a BLAST search to identify the COI gene of the rodent species with the closest identity (http://www.ceropath.org/barcoding_tool/rodentsea).

Inbred CAROLI/EiJ and SPRET/EiJ DNAs were similarly prepared from tissues purchased from the Jackson Laboratory. Initial genotyping was performed by PCR using primers Chr6F (5’ CAAGAGTCCTATGTGTACCTTC 3’) and Chr6Rev (5’ GCAGGCCAATCATAGCACTG 3’) or Chr4F (5’ CAGCAACCACATGGTGACTC 3’) carried out in 50 µl reactions containing 2.5 U of Pfu ultra, 100 ng of template, 0.2 mM dNTPs and 0.5 µM each of the forward and reverse primer. The reaction was performed in a thermal cycler at 95°C for 2 minutes followed by 25 cycles of 95°C for 1 minute, 57°C for 2 minutes and 72°C for 3 minutes.

### Fv1/7 cloning

*Fv1* and *Fv7* were cloned using Q5 high fidelity polymerase (New England BioLabs) with primers Fv1GenStopRev (5’ CCTCCTGATTTTAAGCTCTTTAAC 3’) and either Chr4Fv1 (5’ CCAATTGACAGTGCCAGGACGCC 3’) or Chr6Fv7 (5’ CAGAAGCTCTGTCTTAGGGGAC 3’) to amplify *Fv1* and *Fv7* respectively. The bands were excised from a 1% agarose gel and purified with QIAquick Gel Extraction kit before cloning into the Zero Blunt Topo vector (Invitrogen). Eight colonies from each reaction were picked for sequencing and a FASTA file of the novel sequences is included as Supplementary Information.

Variants were amplified from this vector using Q5 high fidelity polymerase with primers GibsonFv1F (5’ GCCCCCATATGGCCATATGAGATCTGGACGCAGCAGCCGAGTT 3’) and GibsonFv1Rev (5’ATCCCGGGCCCGCGGTACCGAGATCTCCTCCTGATTTTAAGCTCTTTAACTGTTGC 3’), and purified on 1% agarose gels before cloning into a BglII and SalI digested delivery vector using HiFi assembly (New England BioLabs), for use in restriction assays.

### Site directed mutagenesis

A PCR based strategy was used to introduce site directed changes to the *Fv1* or *Fv7* genes. 10 ng of plasmid carrying the gene was used together with 150 ng of each primer containing the altered sequence and spanning the site to be mutated. The reaction was performed using PfuUltra (Agilent) with 18 cycles of denaturation at 95°C for 30 seconds, 55°C for 1 minute and 68°C for 9 minutes 30 seconds. The reaction mixture was then digested with DpnI (New England BioLabs) for 1 hour before using 4 µl for the transformation of XL10 gold ultracompetent cells (Agilent). Colonies were screened for the mutation and verified by sequencing.

### Cells and virus production

MDTF and 293T cells were maintained in Dulbecco’s modified Eagle’s media containing 10% fetal calf serum and 1% penicillin/streptomycin. Viruses were made by transient transfection of 293T cells as described previously (17, 53, 54). To make delivery viruses for transducing permissive MDTF with *Fv1/Fv7*, pczVSVG and pHIT60 were co-transfected with pLIEYFP carrying the *Fv1* or *Fv7* variant. Apart from the foamy viruses, the tester viruses were all pseudotyped with VSVG. N-tropic, B-tropic, Mo-MLV and NR-tropic MLV were made by co-transfecting pczVSVG and pfEGFPf with either pCIGN, pCIGB, pHIT60 or pCIGN(L117H) respectively. EIAV was made by co-transfection of pczVSVG, pONY3.1 and pONY8.4ZCG (55) while FIV was produced with pczVSVG, pFP93 and pGiNWF-G230 (56). PFV and FFV were generated using pciSFV-1envwt and either pczDWP001 or pcDWF003 respectively (57). MLVs and FIV were aliquoted and frozen at -80°C after harvesting while EIAV and foamy viruses were used fresh. Transduction using EIAV and FIV were performed in the presence of 10 µg/ml polybrene.

### Restriction assay

Restriction activity was measured using a flow cytometry-based assay as described previously (53, 54). Briefly, the *Fv1/Fv7* genes were delivered into permissive MDTF cells using a Mo-MLV-based bi-cistronic vector which also contains EYFP in the same transcriptional unit so that all cells that express the restriction factor would also fluoresce yellow. Three days later, the cells were challenged with a tester virus that carried EGFP so that infected cells fluoresced green. Three days post-infection, the cells were analyzed by flow cytometry to obtain the ratio of the number of infected cells (green) containing restriction factors (yellow) to infected cells that did not contain restriction factors (non-yellow). A ratio of less than 0.3 was indicative of restriction while that which was greater than 0.7 showed the absence of restriction. A value between 0.3 and 0.7 was taken to represent partial restriction.

In order to study the effect of expressing two different restriction factors in the same cell, the assay described above was modified by transducing MDTF (factors expressed from retroviral promoter) or R18 cells (factors expressed from inducible promoter) with the EYFP vector containing the first restriction gene together with a vector containing the second restriction gene and mScarlet. Three days later, the cells were challenged with a tester virus that carried EGFP. Cells containing one factor were either yellow or red while those transduced with both factors were yellow and red. The different populations, together with untransduced cells, were analyzed by flow cytometry to obtain the percentage of infected (green) cells in each population.

### Phylogenetic analysis

Nucleotide sequences corresponding to nucleotides 1-1299 of Fv1CAR1, i.e. without the variable tail due to the insertion of SINE elements in *M. cooki* and *M. cervicolor*, were aligned with MAFFT v7.271 (58, 59) and used to build an ML tree with a GTR+CAT model using FastTree v2.1.9 (60).

### Analysis of Fv1 transcription

pGL4.10 (Promega) plasmids were produced with synthesized DNAs representing the region from 150 to 350 nucleotides 5’ of the *Fv1* ATG. Mutated constructs were produced for the putative initiator elements by replacing the sequences with adenine. These, and the control SV40-driven pGL4.13, were introduced to MDTF cells with GeneJuice (Merck) for harvest after 24 hours. 5x10^4^ cells were re-suspended in phenol-free media, mixed with Bright-Glo luciferin (Promega) and assayed according to the manufacturer’s instructions using opaque-walled black 96 well plates. 4 separate experiments were assayed in triplicate.

### Prediction of INR elements

The sequence preceding the Fv1 CDS was scanned with a predefined PWM (41) using inbuilt functionality within UGENE v1.28 (61).

## Acknowledgements

This work was supported by the Francis Crick Institute, which receives its core funding from Cancer Research UK (FC010162), the UK Medical Research Council (FC010162) and the Wellcome Trust (FC010162). JPS holds an Investigator Award from the Wellcome Trust (108012/Z/15/Z). SM was supported by the French ANR CERoPath (ANR 07 BDIV 012) and FutureHealthSEA (ANR-17-CE35-0003-02).

## Supporting information

**Fig S1.**
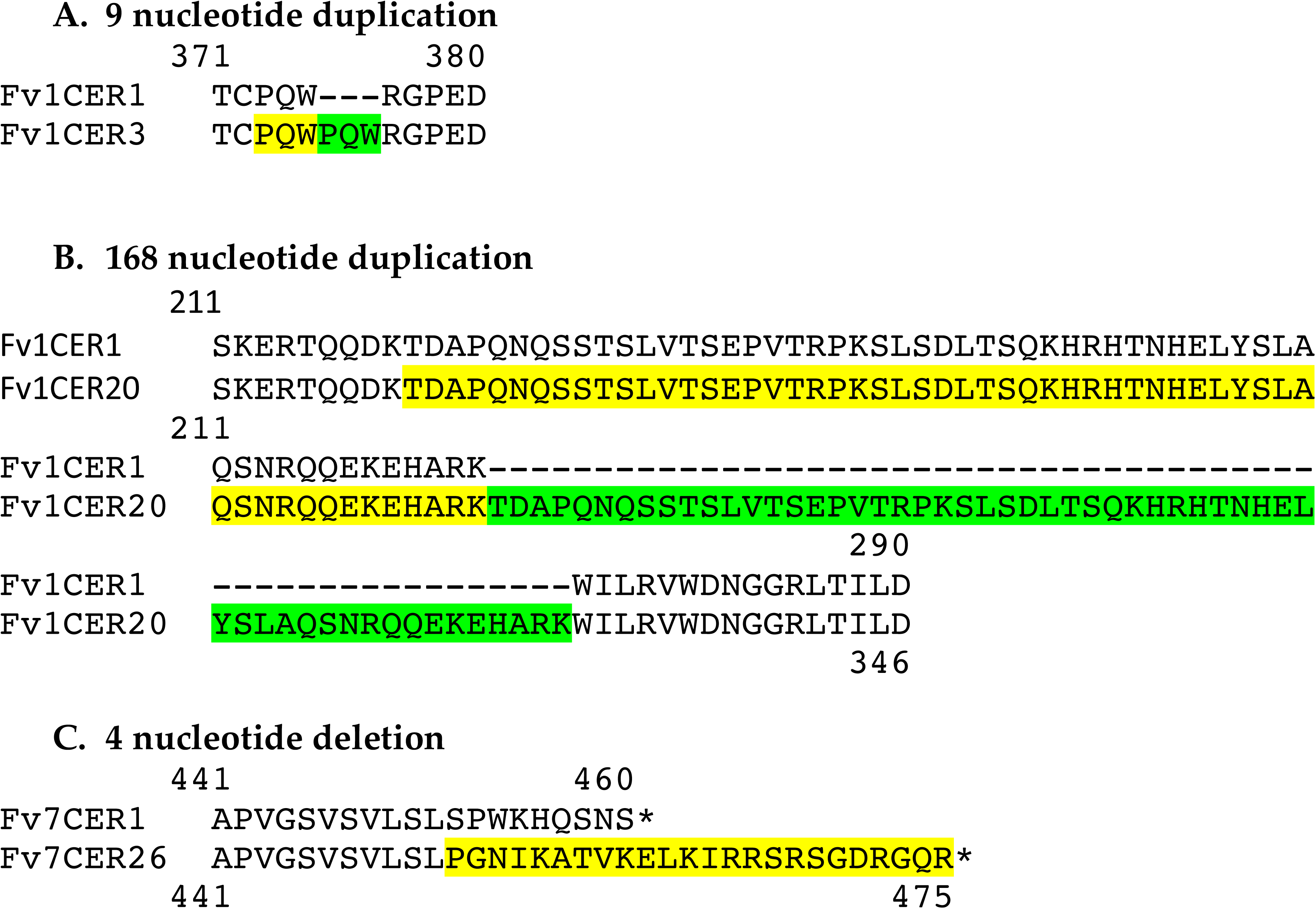
Some effects of duplication or deletion on predicted Fv1/Fv7 sequences. **A.** Alignment of the amino acid sequences of Fv1CER1 and Fv1CER3. The target sequence of proline, glutamine and tryptophan (PQW) in Fv1CER3 is boxed in yellow while the repeat is shown in green. **B**. Alignment of the amino acid sequences of Fv1CER1 and Fv1CER20. The target sequence of Fv1CER20 is boxed in yellow while the repeat is shown in green. **C.** Alignment of the amino acid sequence at the C-terminus of Fv7CER1 and Fv7CER26 showing the extension of the C-terminus of Fv7CER26 due to frame-shifting following the deletion of 4 nucleotides. The alternative sequence caused by the frame-shift is shown in yellow.

**Fig S2.**
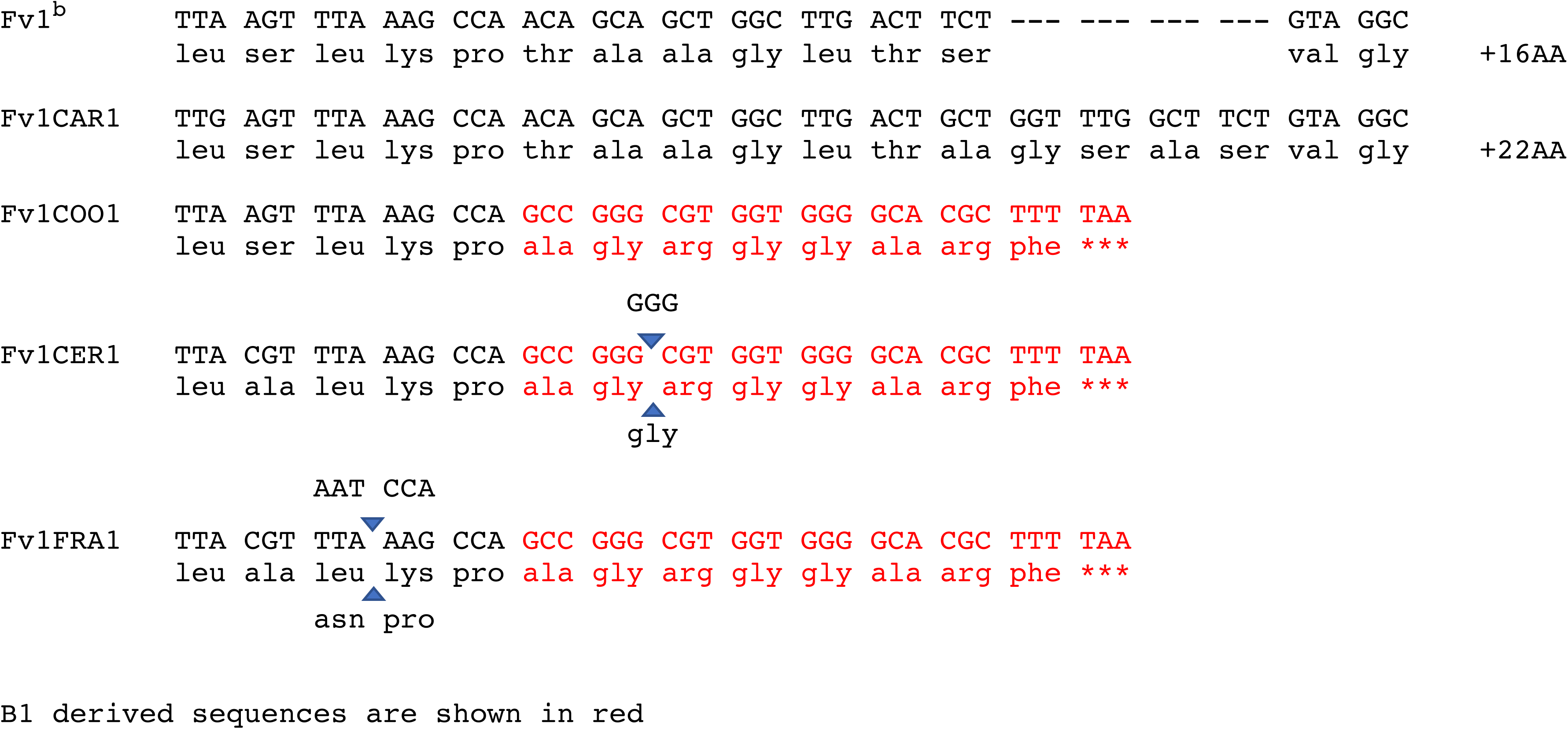
B1 truncation of Fv1 C terminal region. The Fv1 C terminal region from *M. caroli*, *M. cookii*, *M. cervicolor* and *M. fragilicauda* in comparison to *Fv1^b^*. Sequences deriving from B1 repeats are highlighted in red. Indel variation between sequences within each species are indicated by blue arrows and corresponding nucleotides and residues.

**Fig S3.**
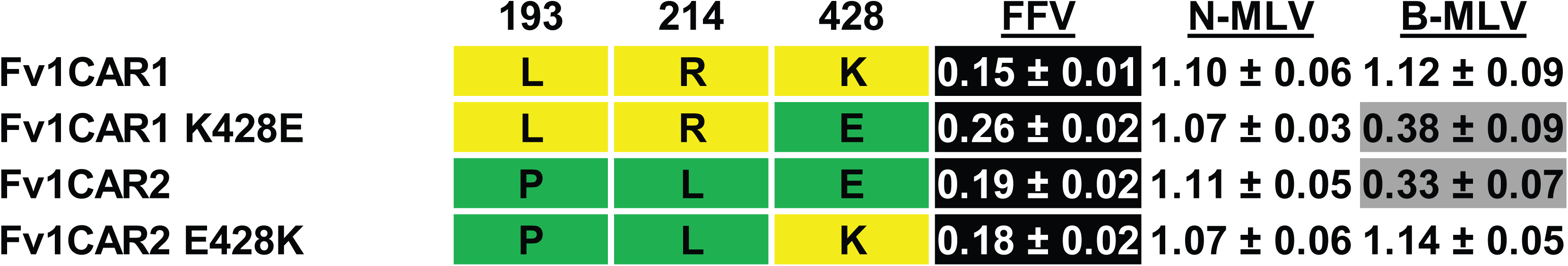
Mapping specificity residues: Fv1CAR1. Residues differing between the restricting and non-restricting variants in the C-terminal region are shown on the left while restriction data are presented on the right of the figure. These variants were introduced into permissive MDTF cells using a retroviral vector also containing the EYFP marker and challenged with EGFP-carrying virus to allow calculation of restriction capacity (see Materials and Methods Values are the means and standard deviations of at least 4 experiments.

**Fig S4.**
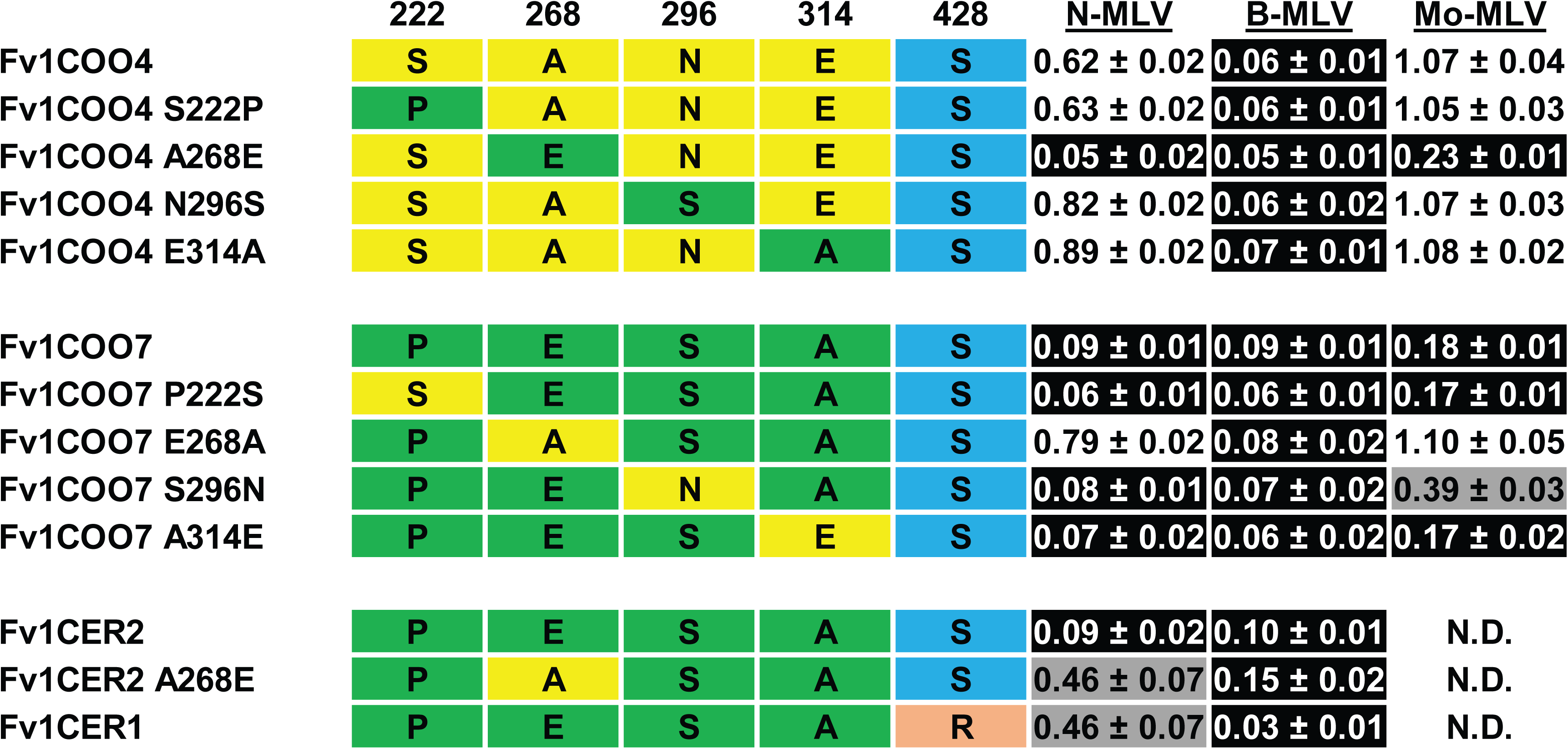
Mapping specificity residues: Fv1COO4. Residues differing between the restricting and non-restricting variants in the C-terminal region are shown on the left while restriction data are presented on the right of the figure. These variants were introduced into permissive MDTF cells using a retroviral vector also containing the EYFP marker and challenged with EGFP-carrying virus to allow calculation of restriction capacity (see Materials and Methods Values are the means and standard deviations of at least 4 experiments.

**Fig S5.**
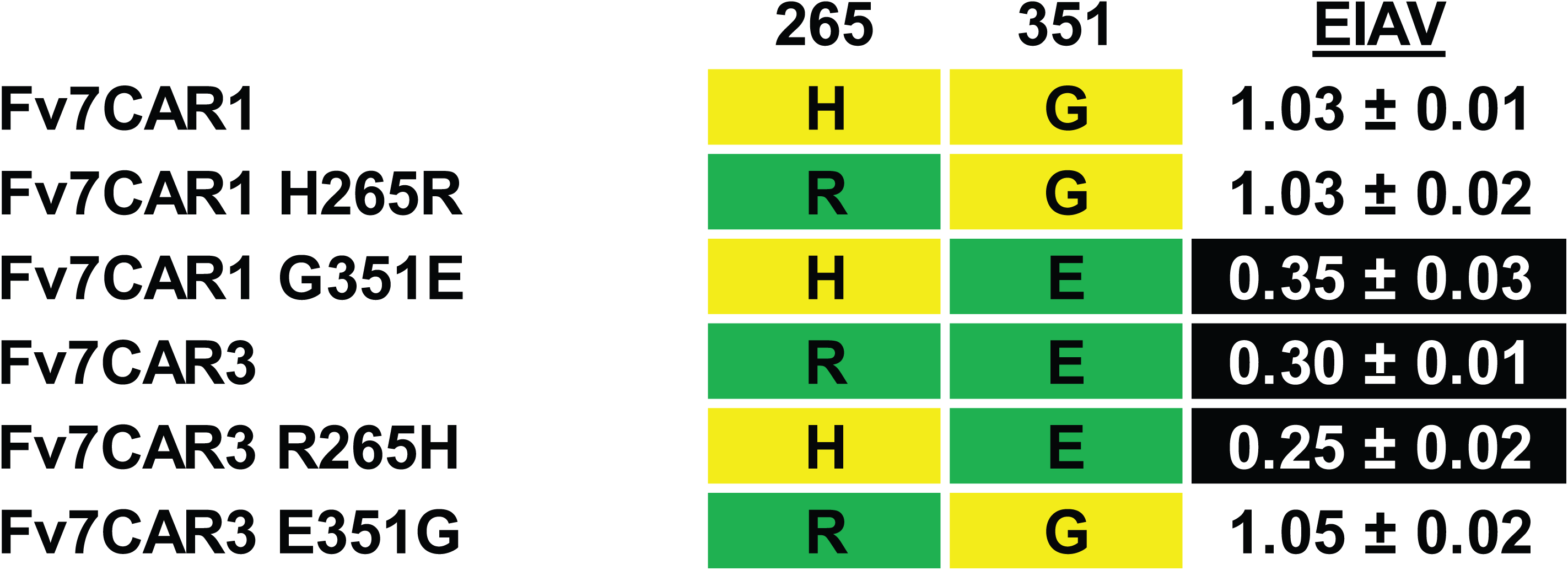
Mapping specificity residues: Fv7CAR1. Residues differing between the restricting and non-restricting variants in the C-terminal region are shown on the left while restriction data are presented on the right of the figure. These variants were introduced into permissive MDTF cells using a retroviral vector also containing the EYFP marker and challenged with EGFP-carrying virus to allow calculation of restriction capacity (see Materials and Methods Values are the means and standard deviations of at least 4 experiments.

**File S1.**
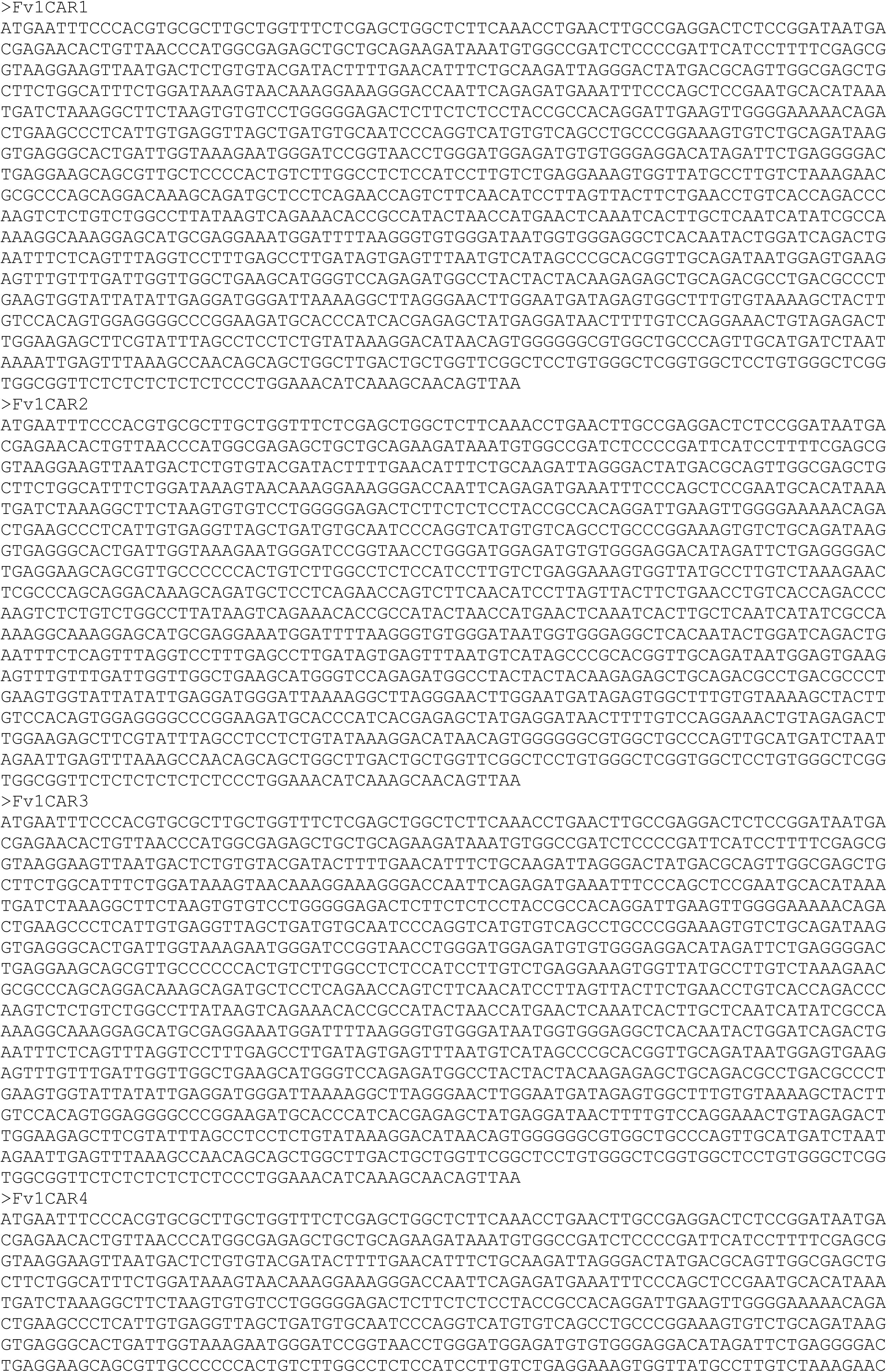
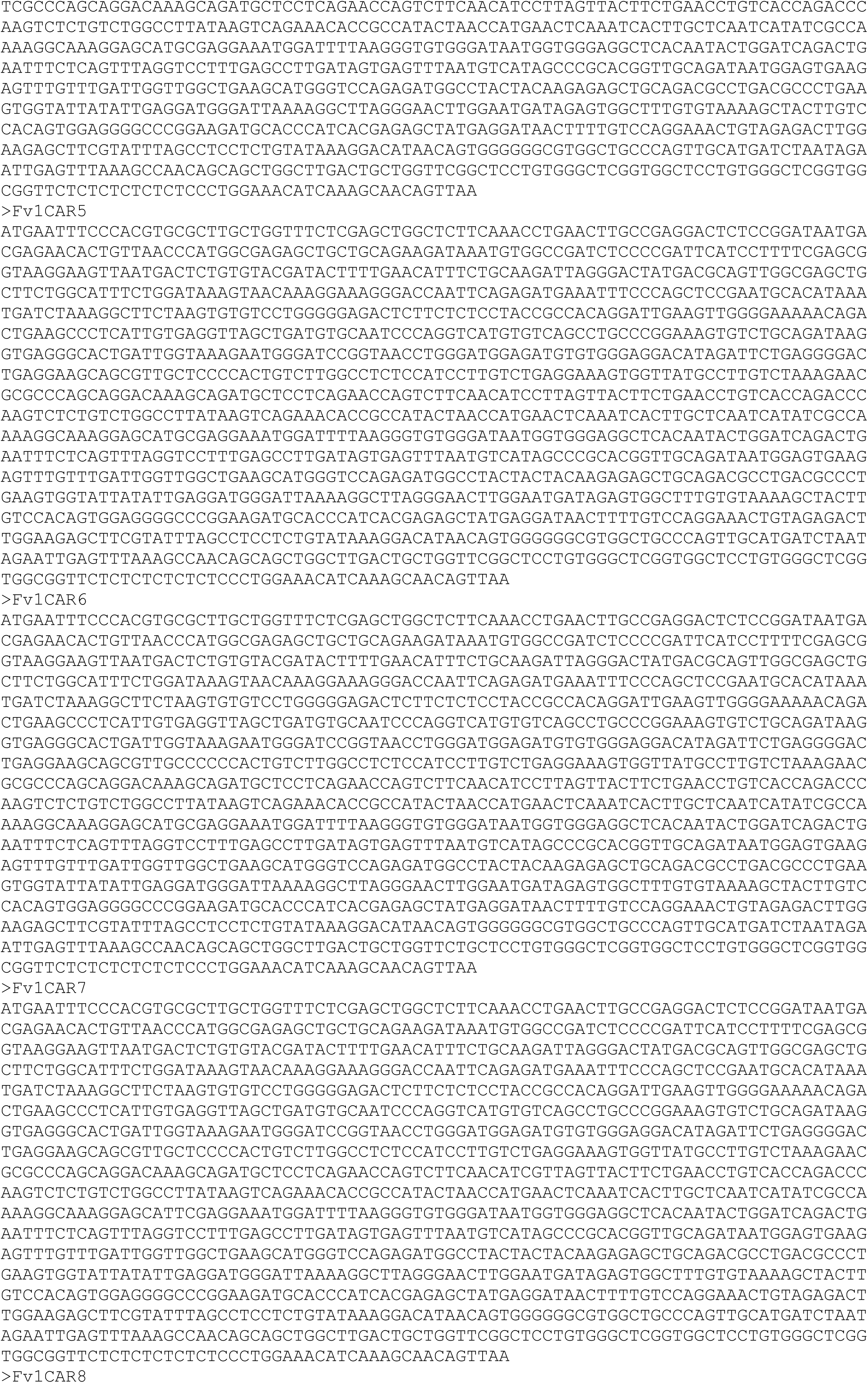
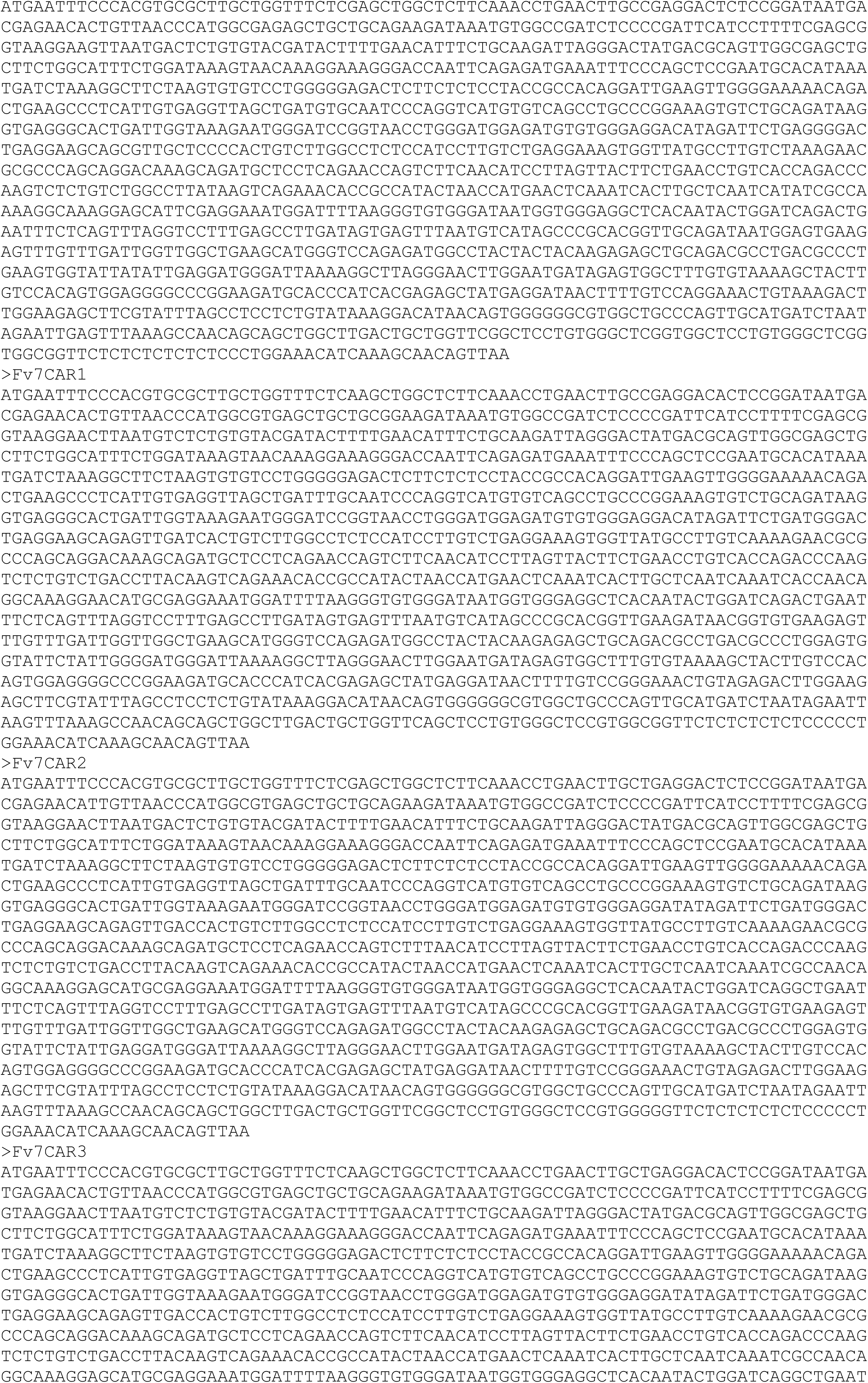
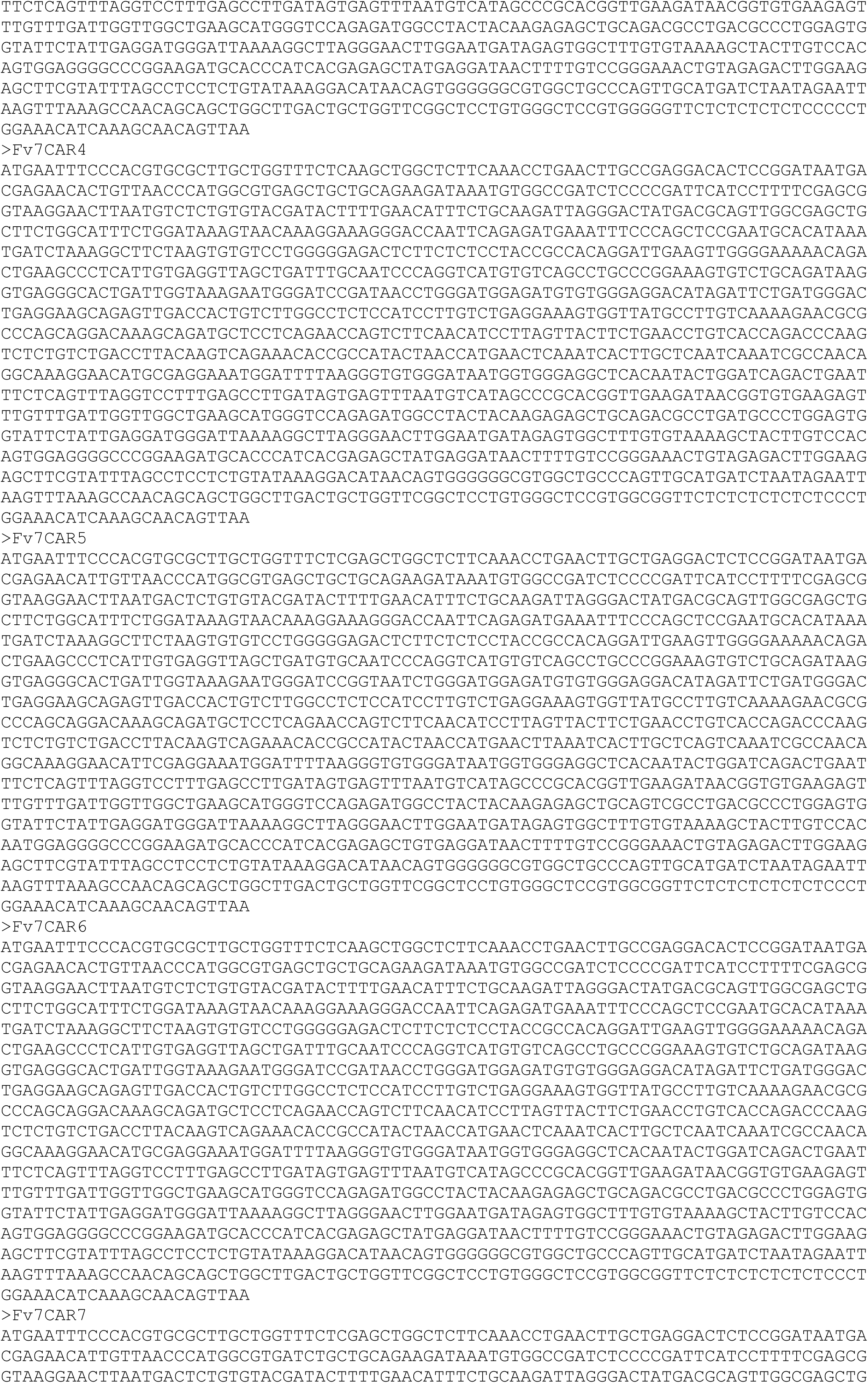
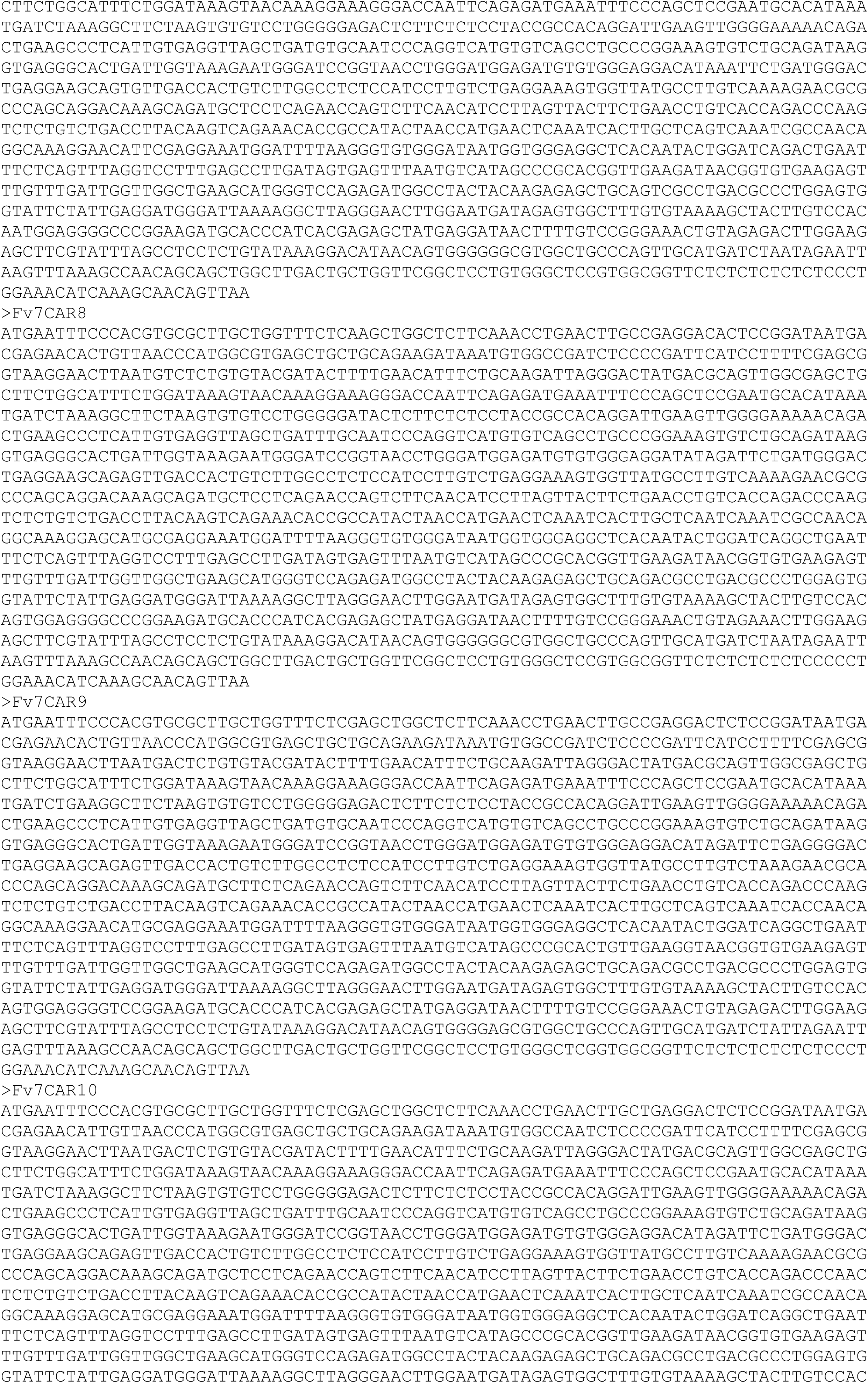
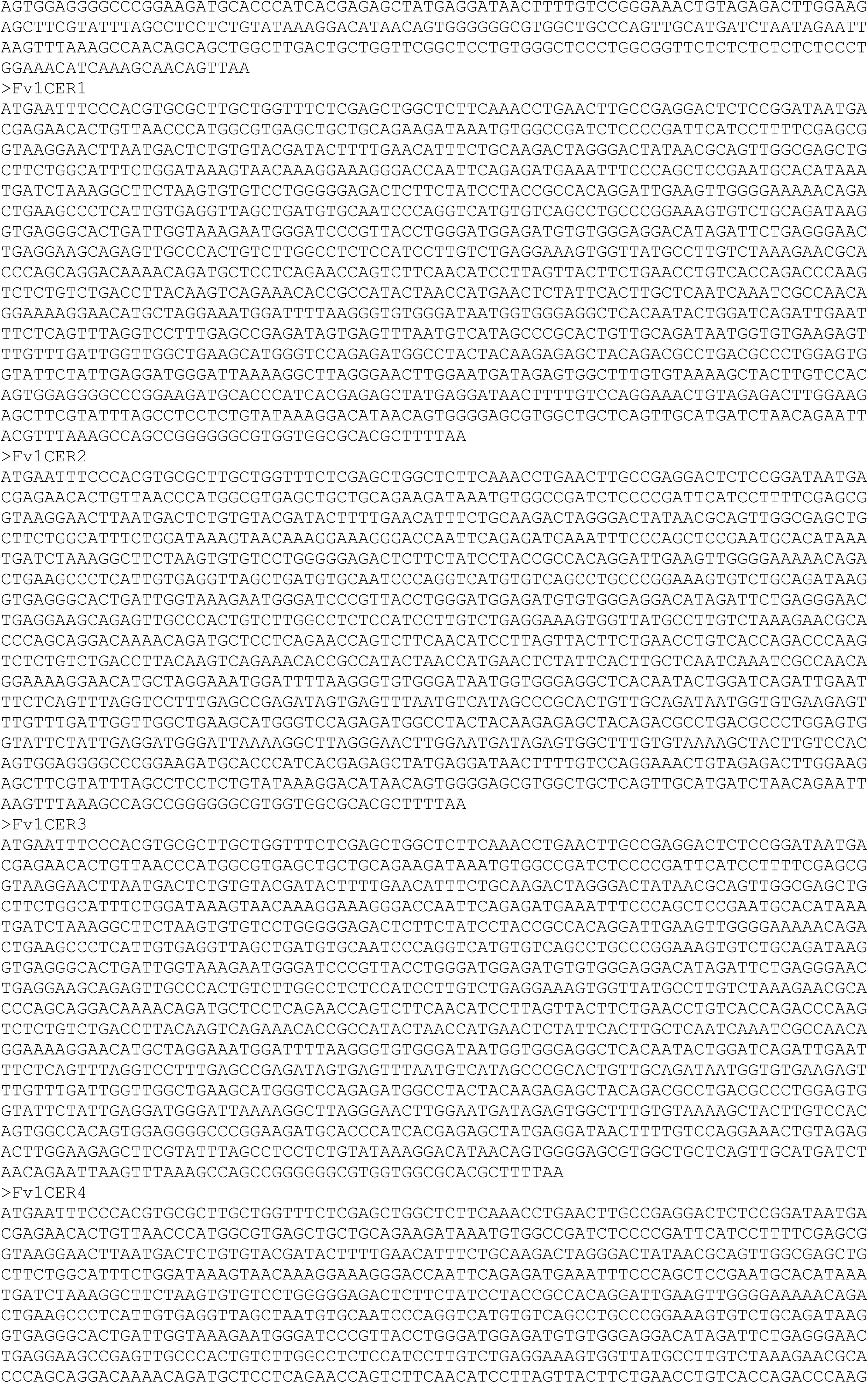
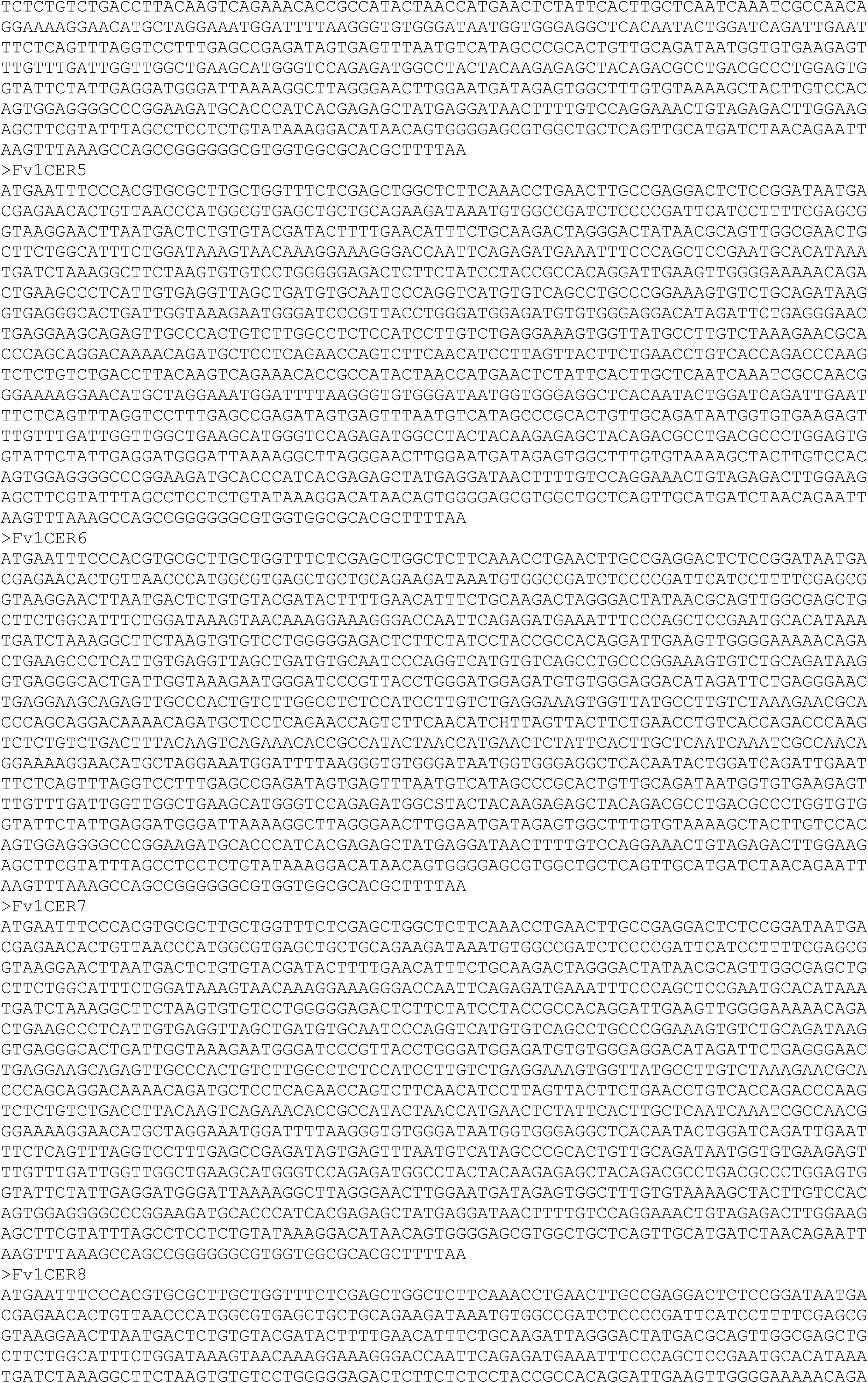
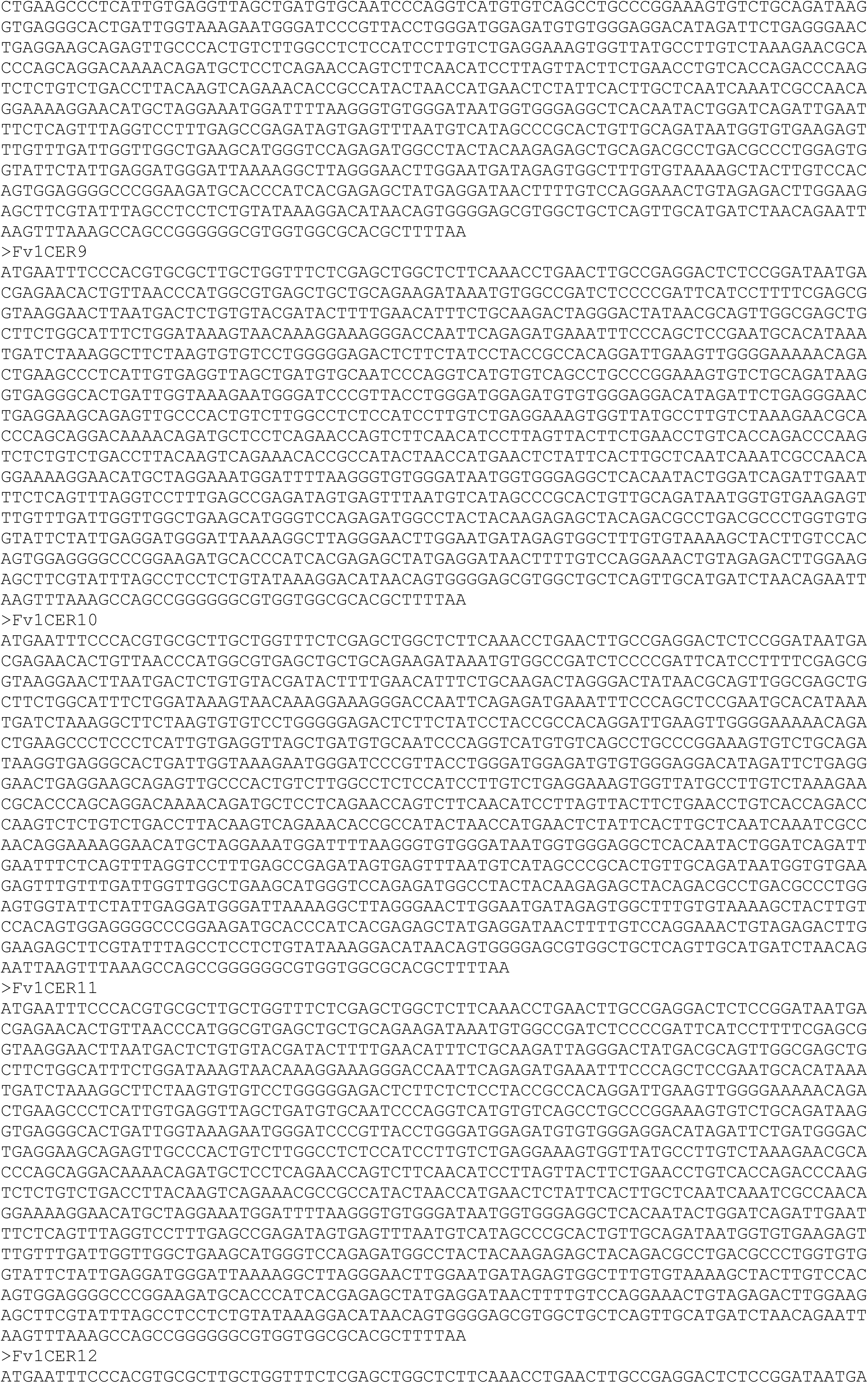
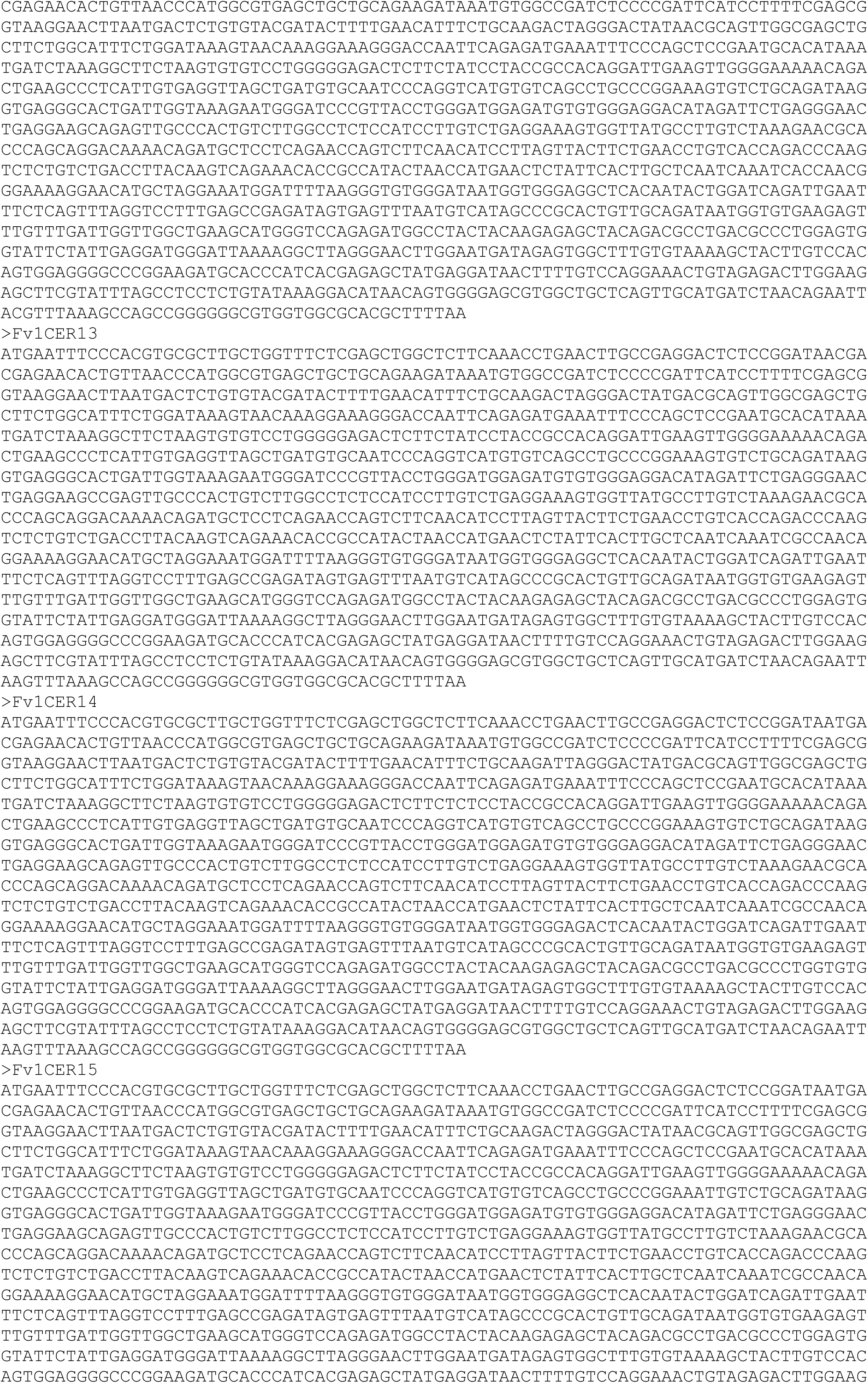
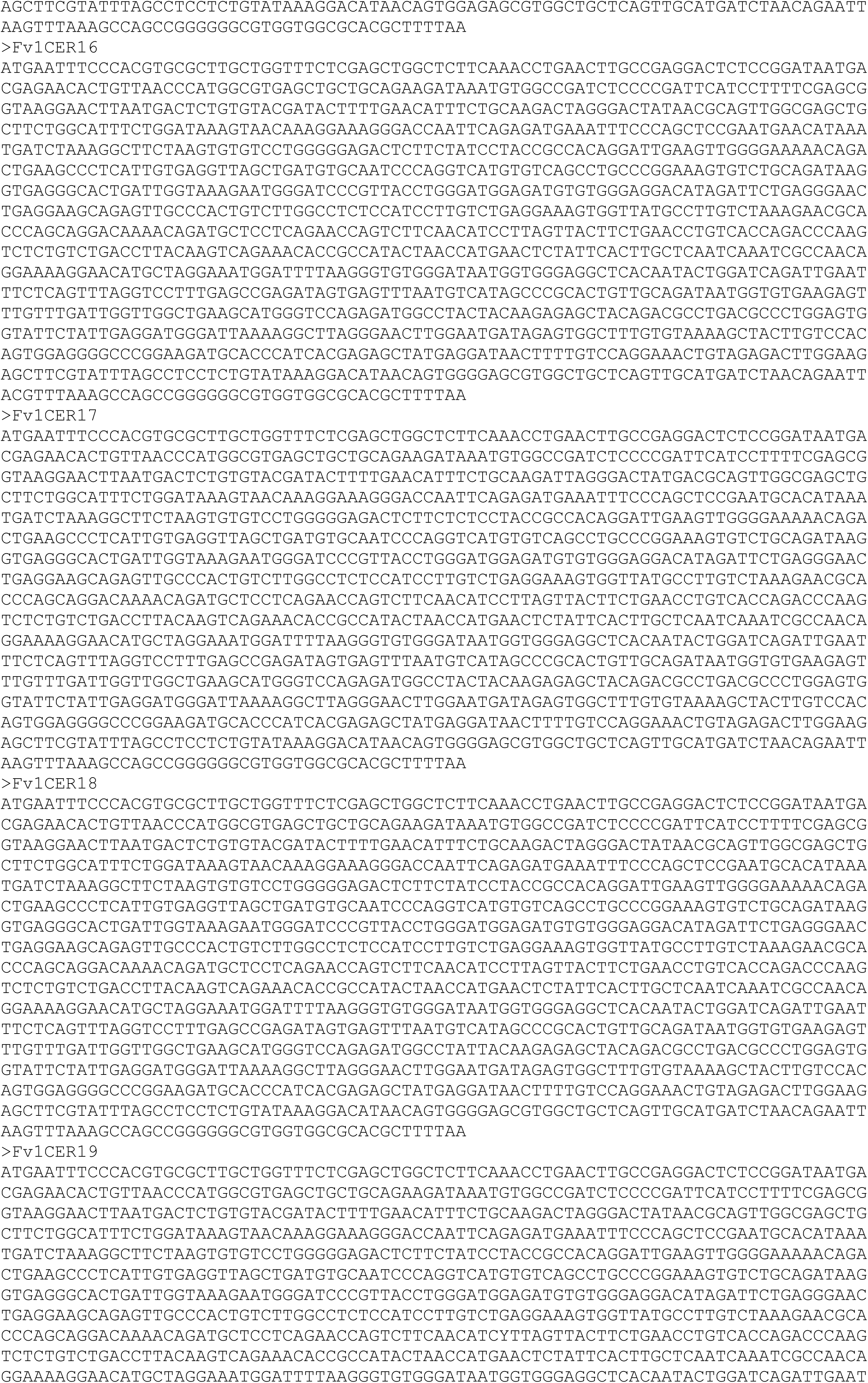
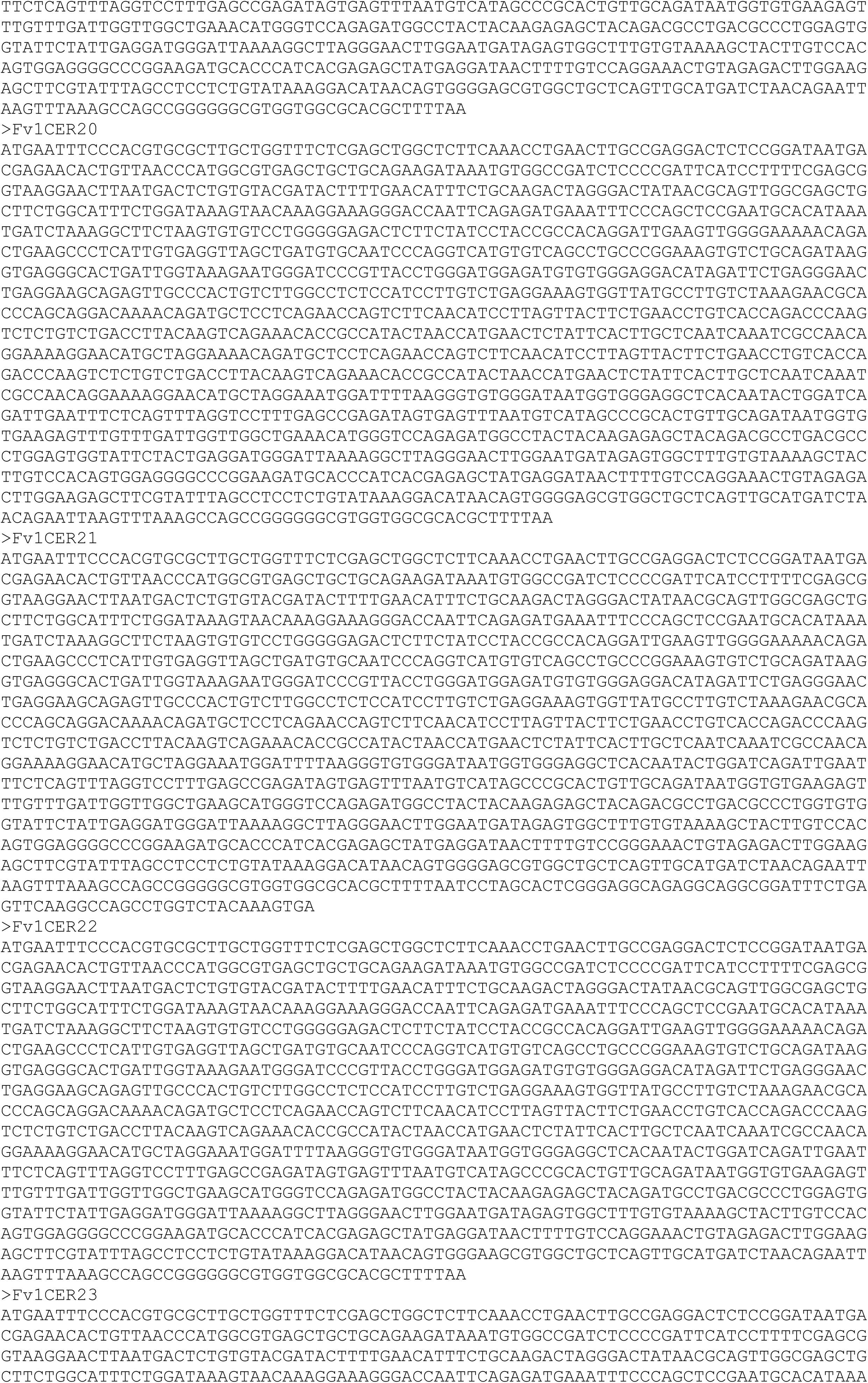
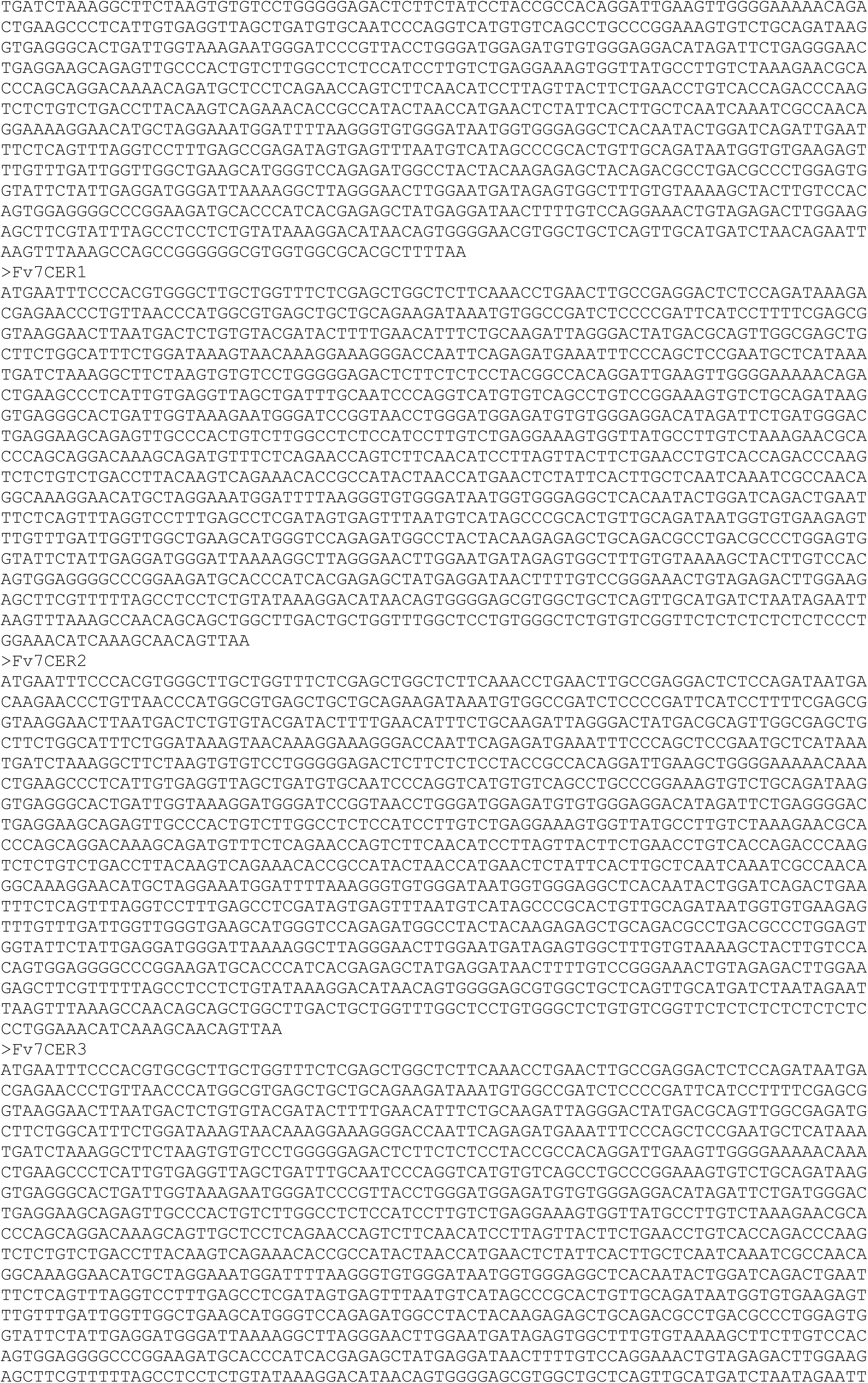
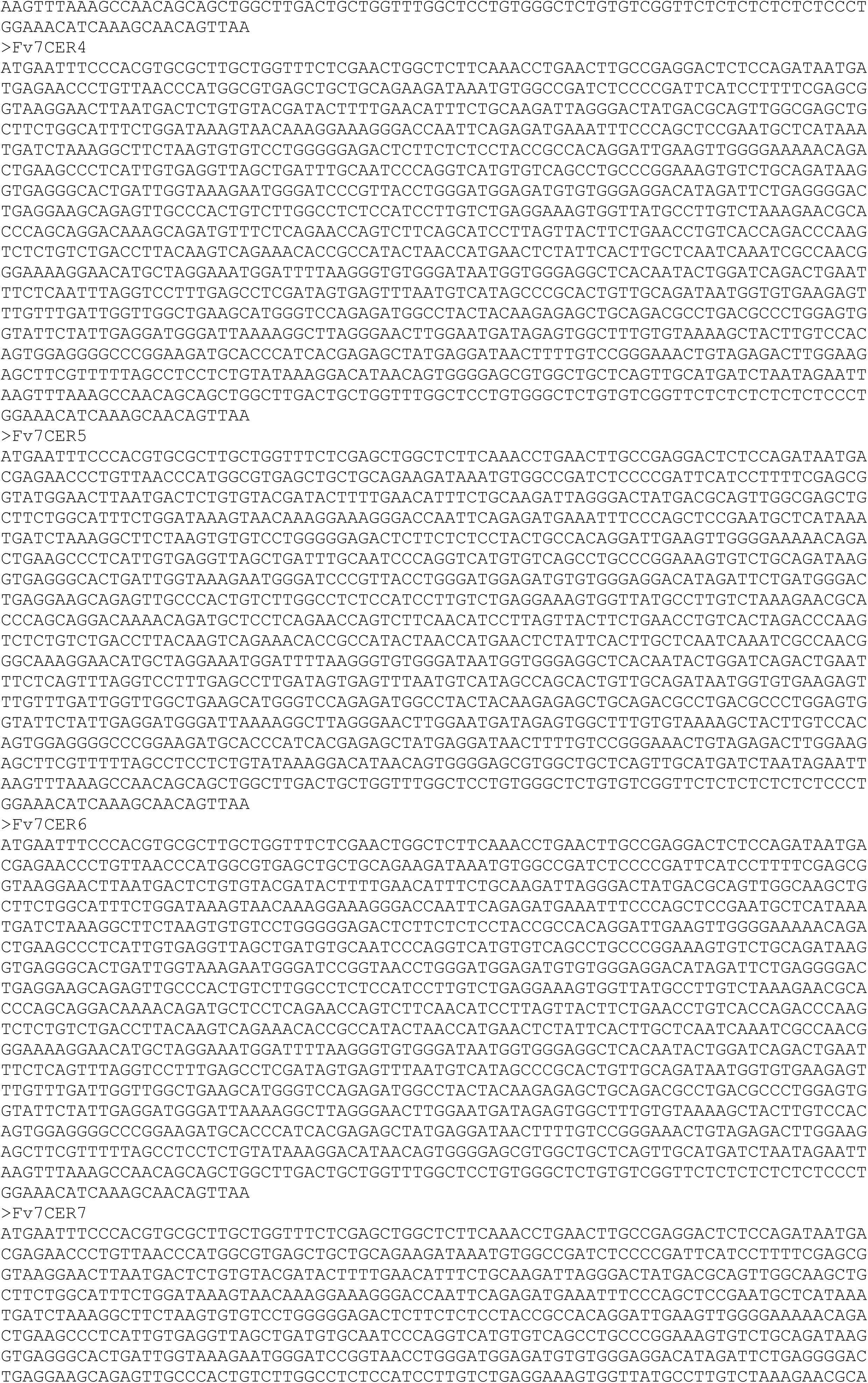
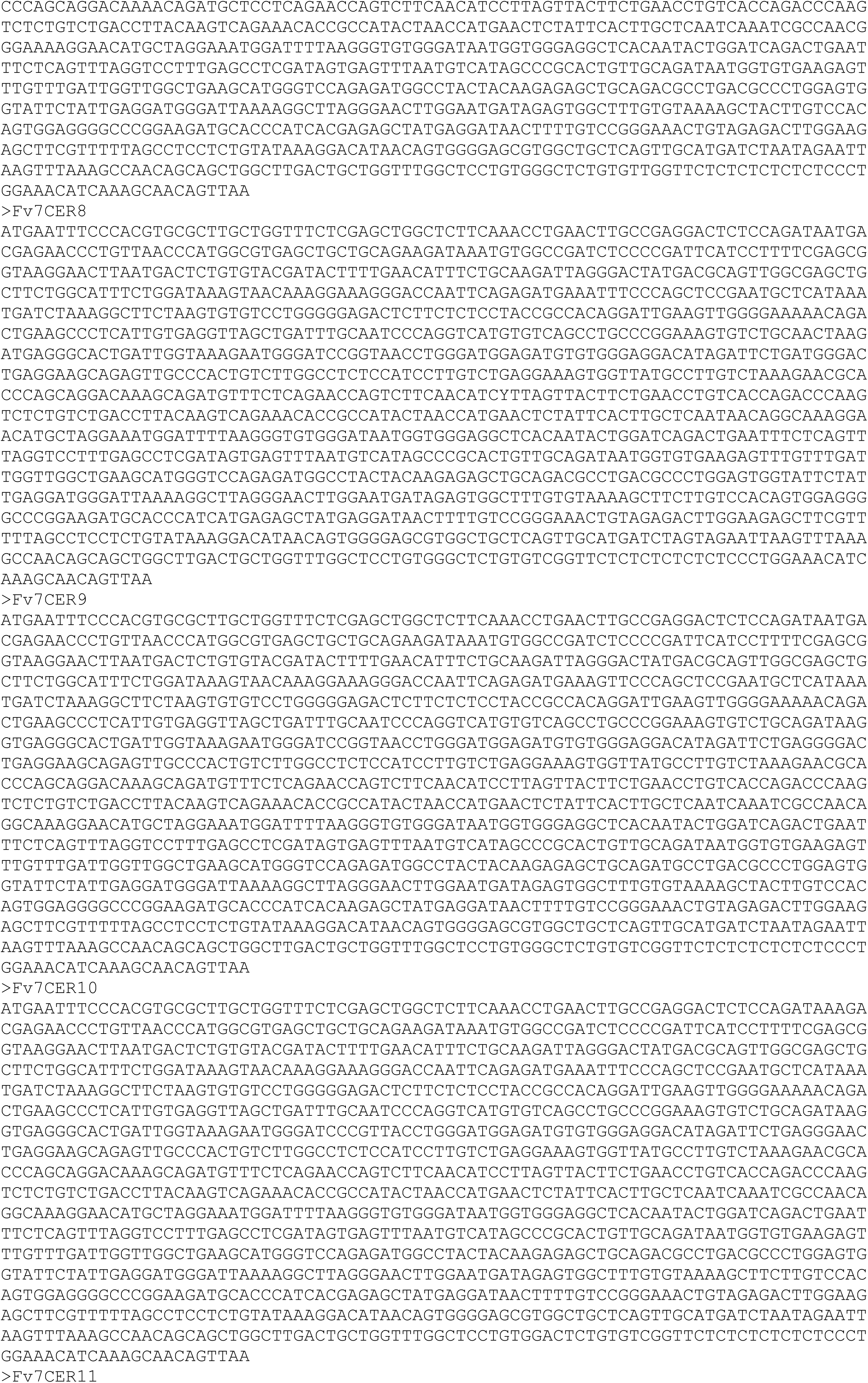
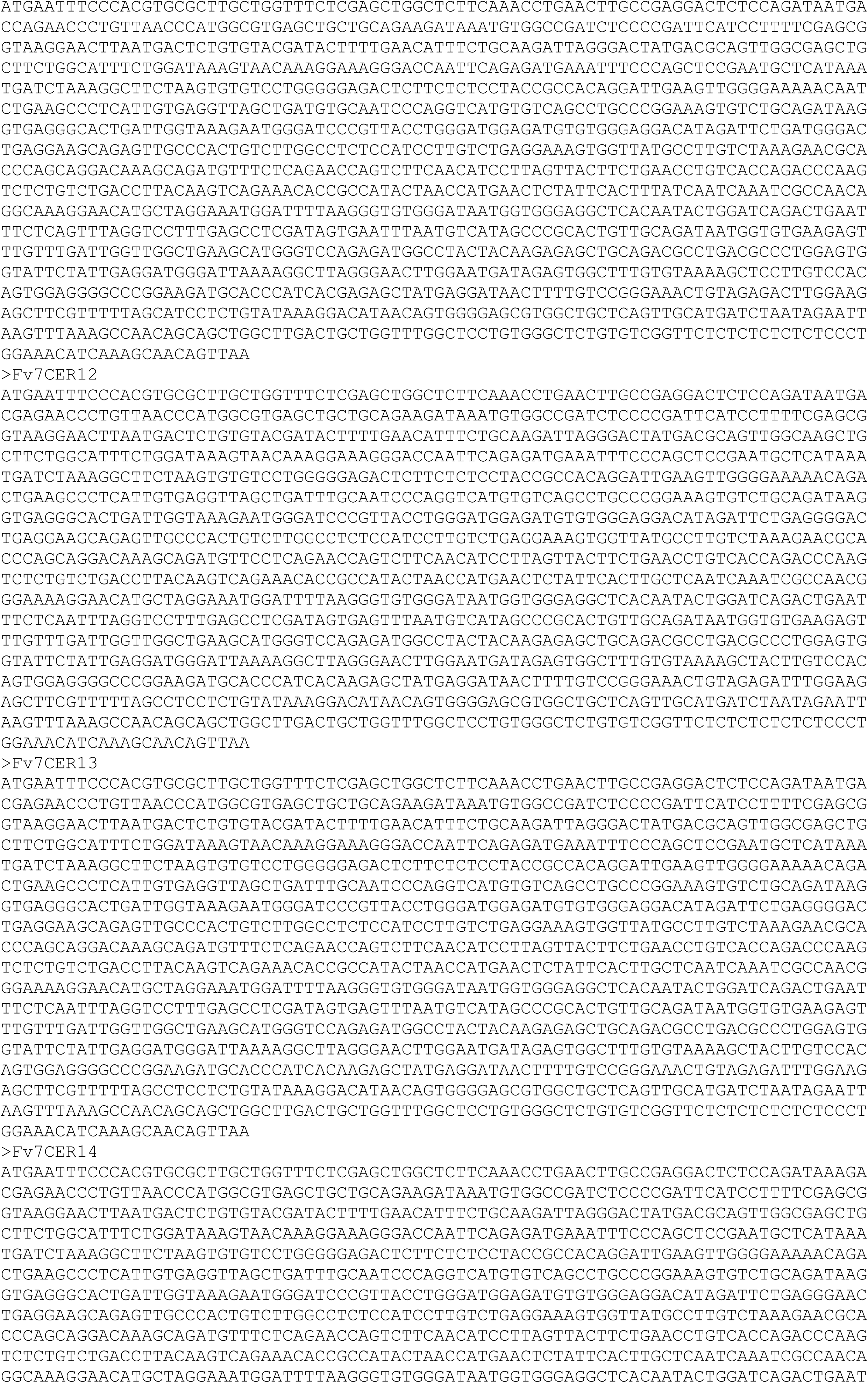
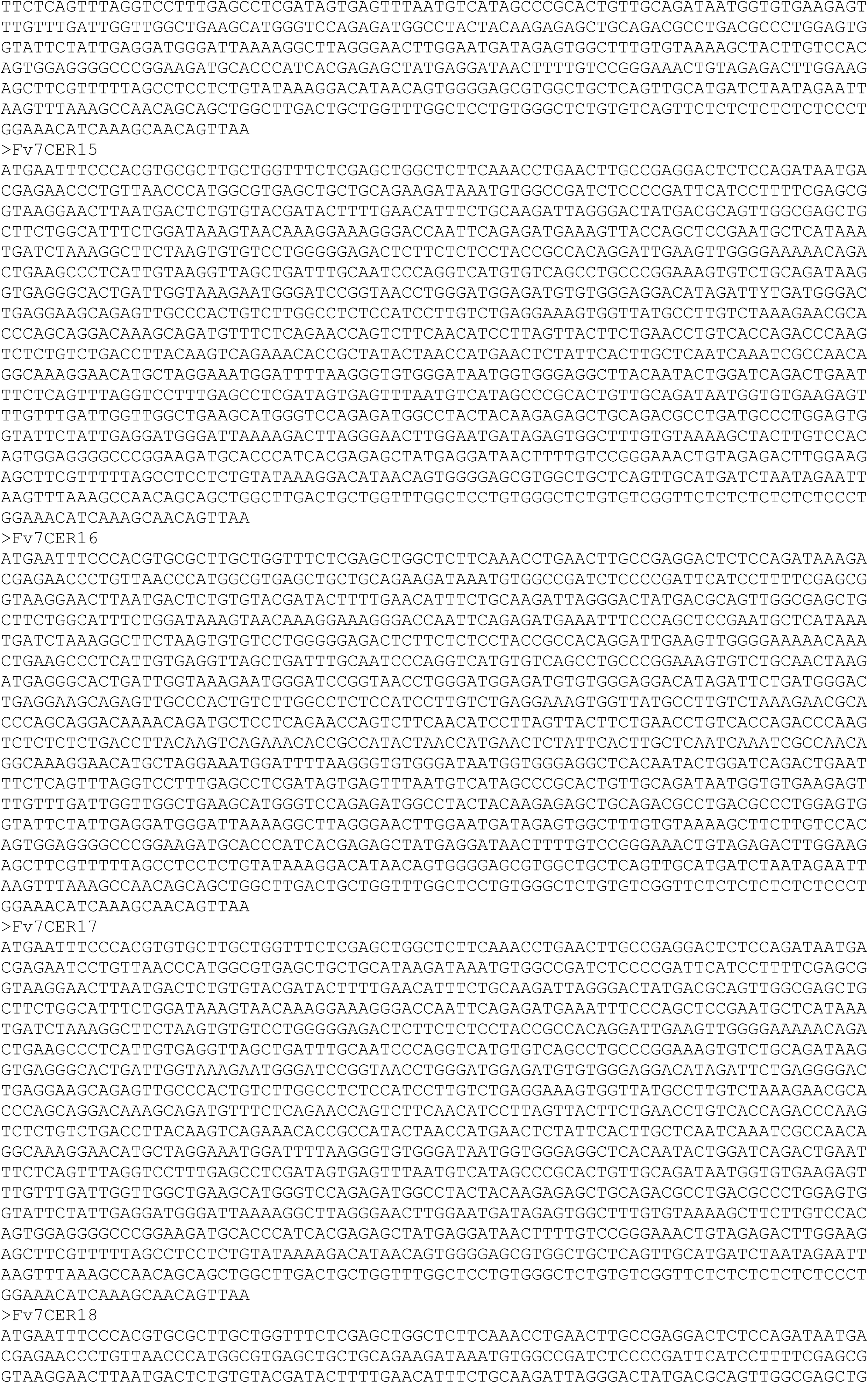
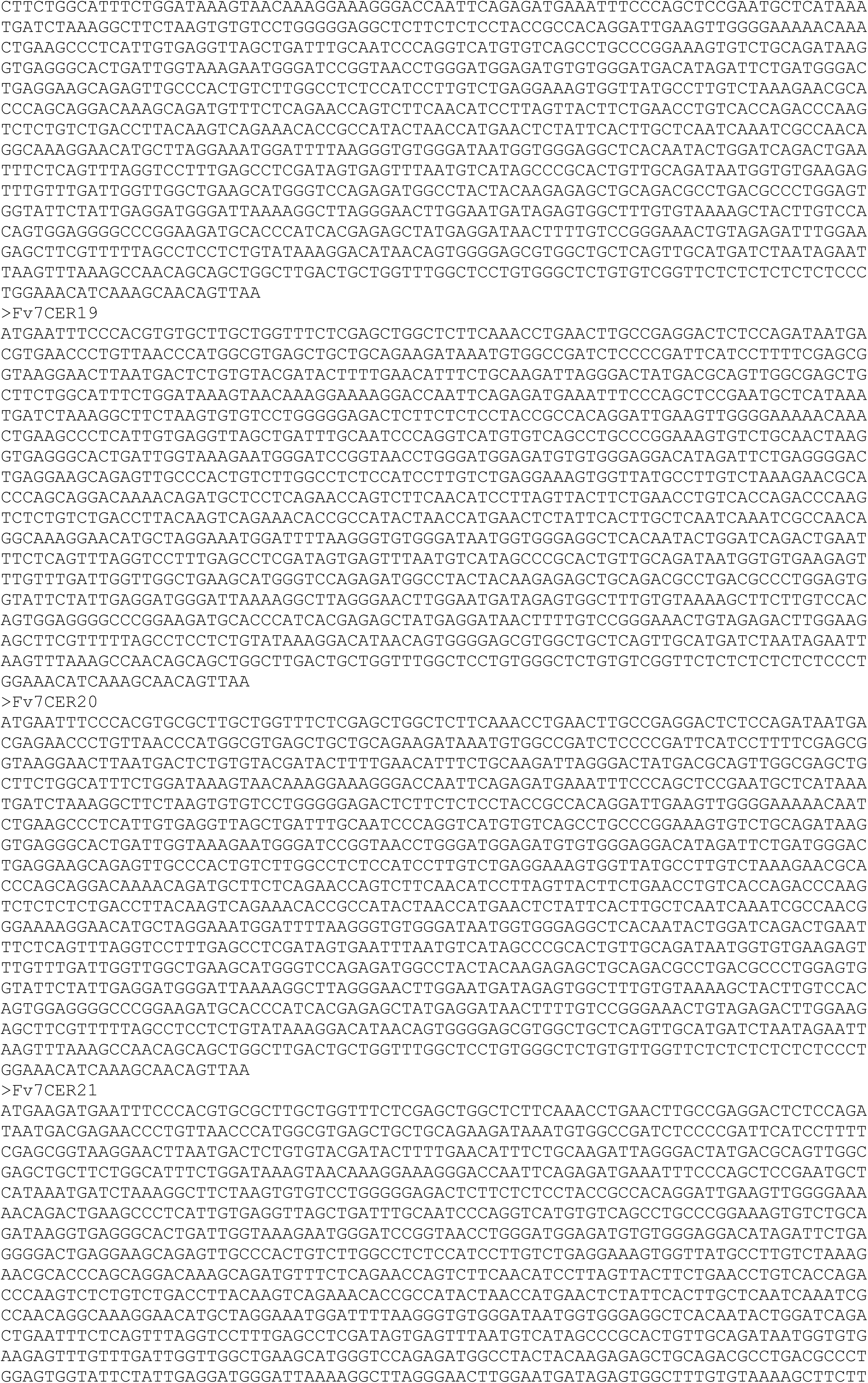
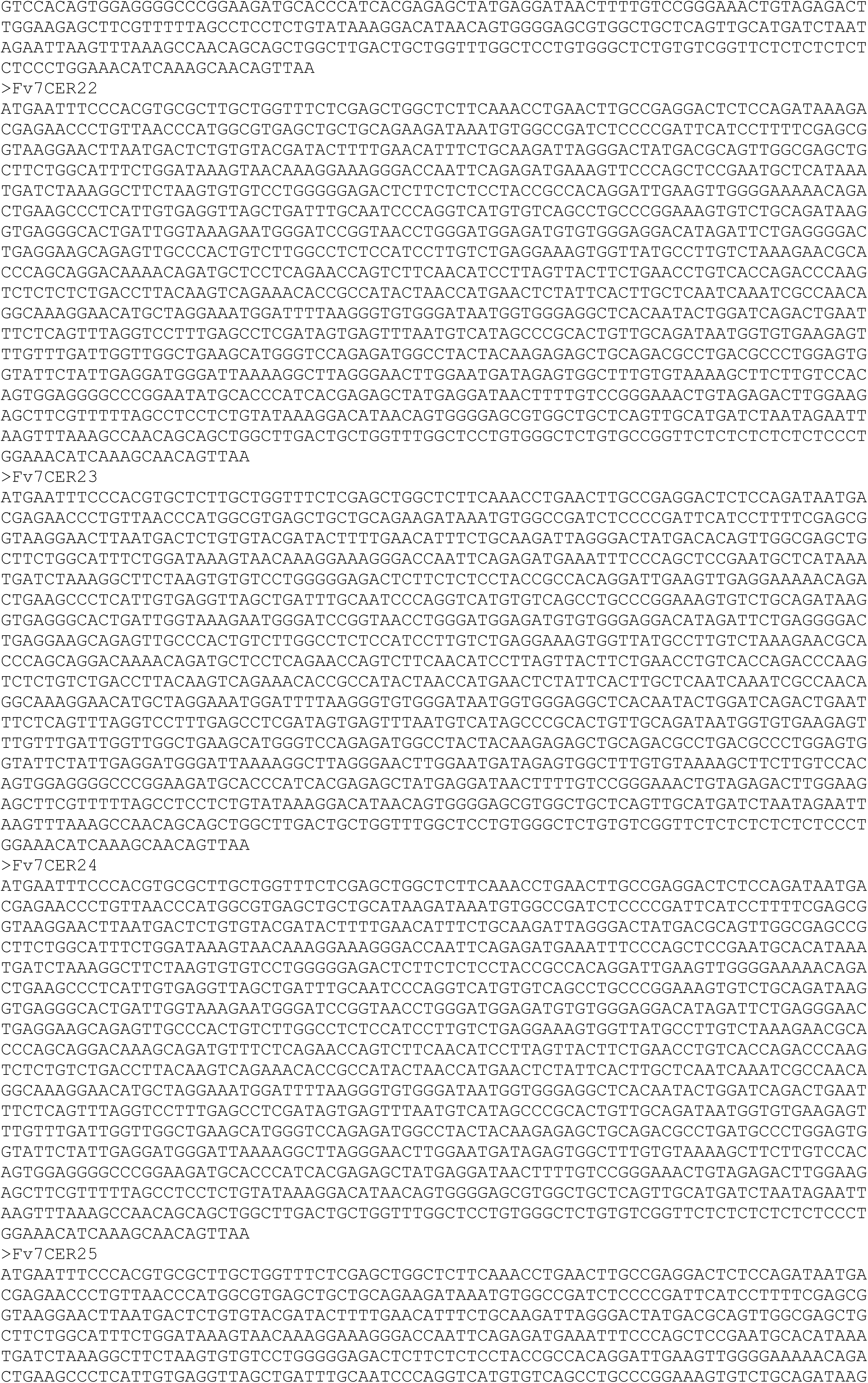
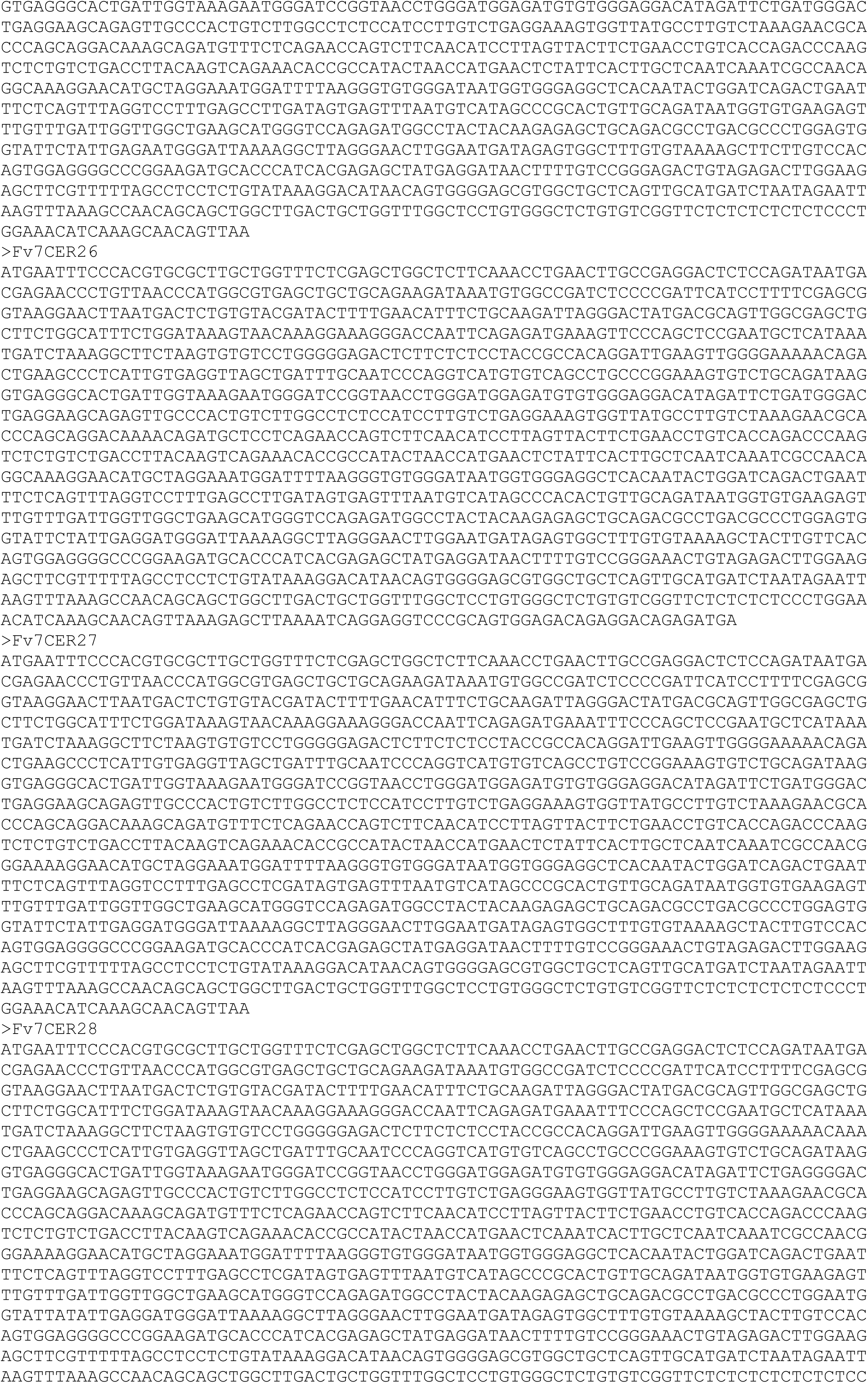
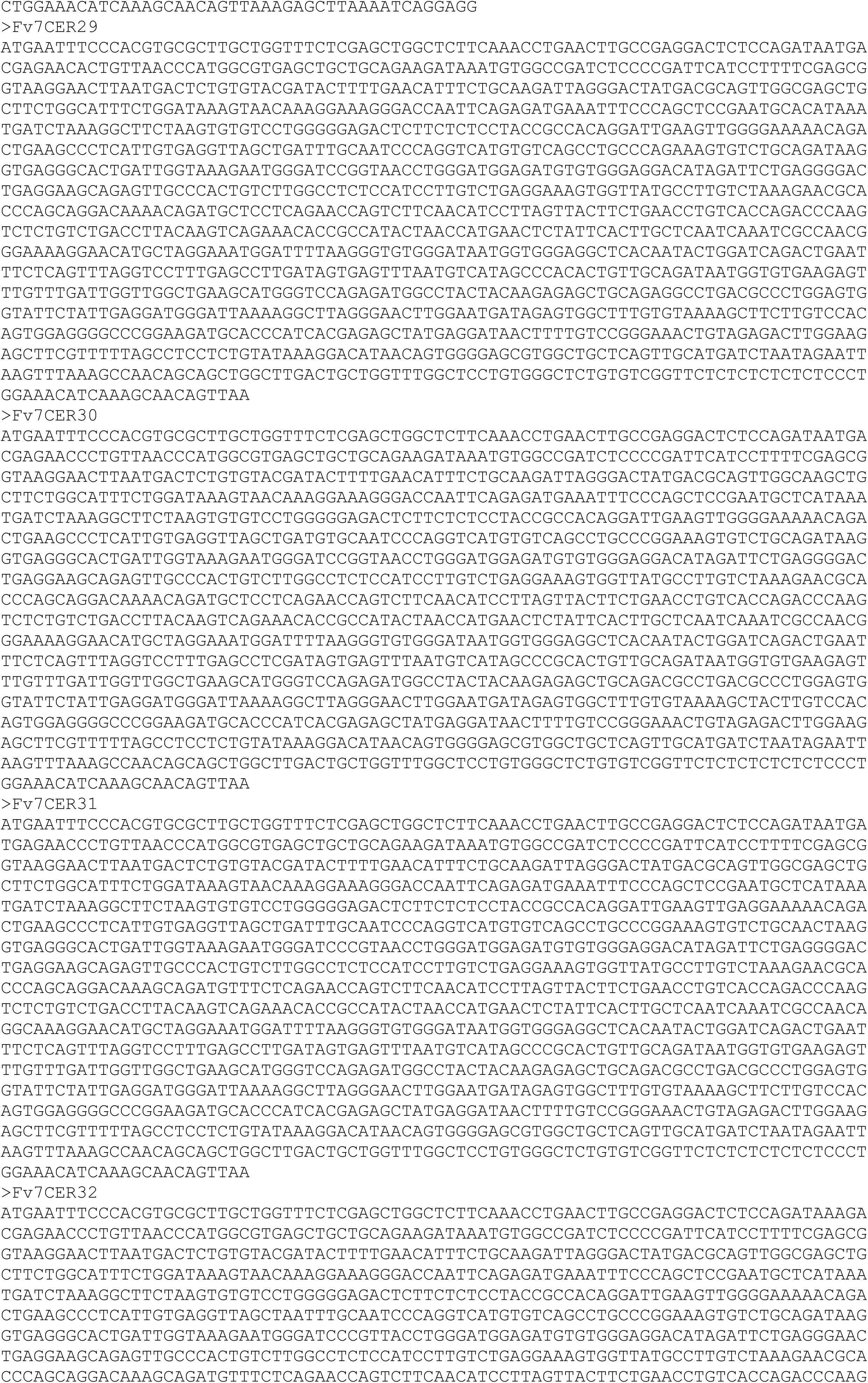
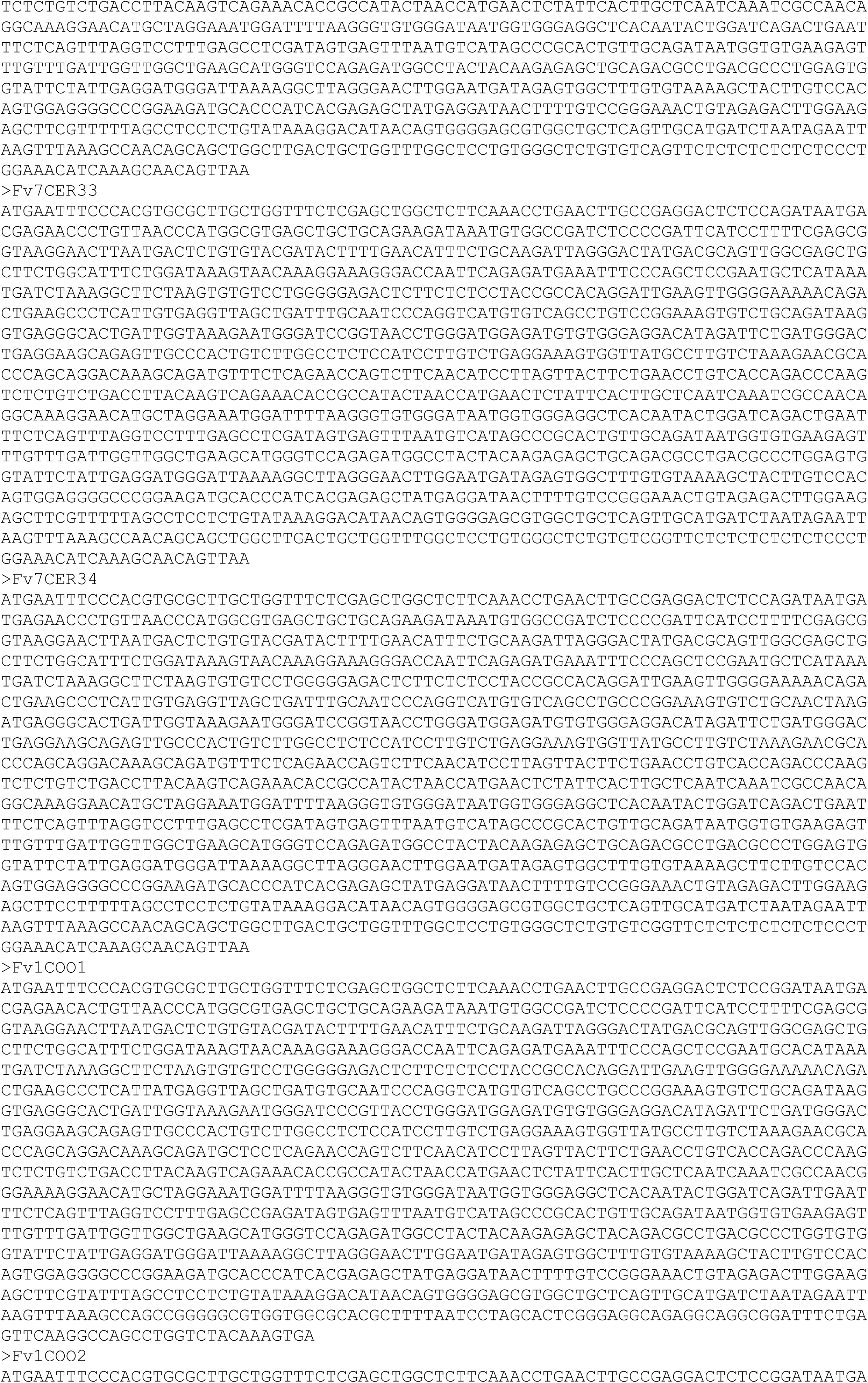
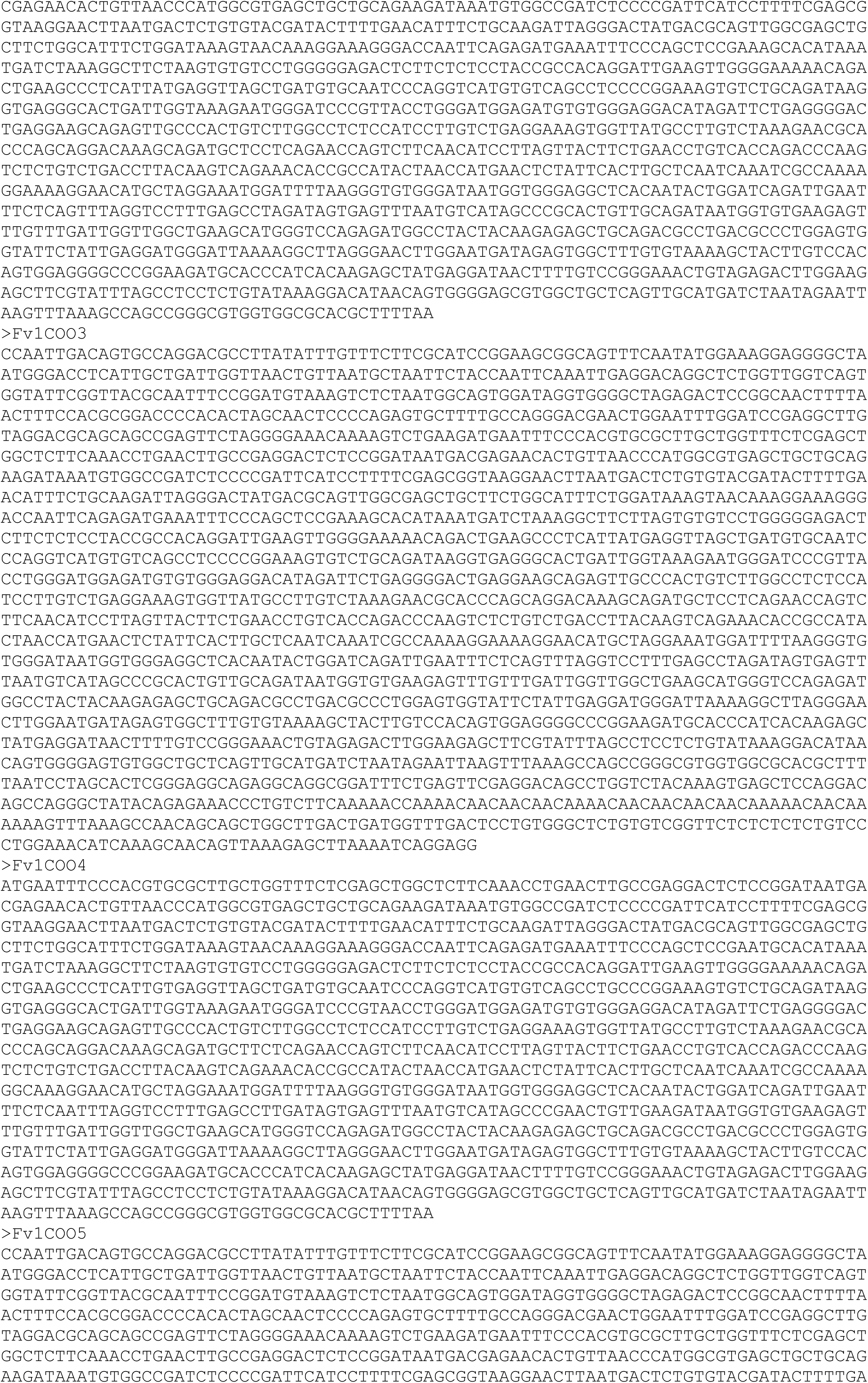
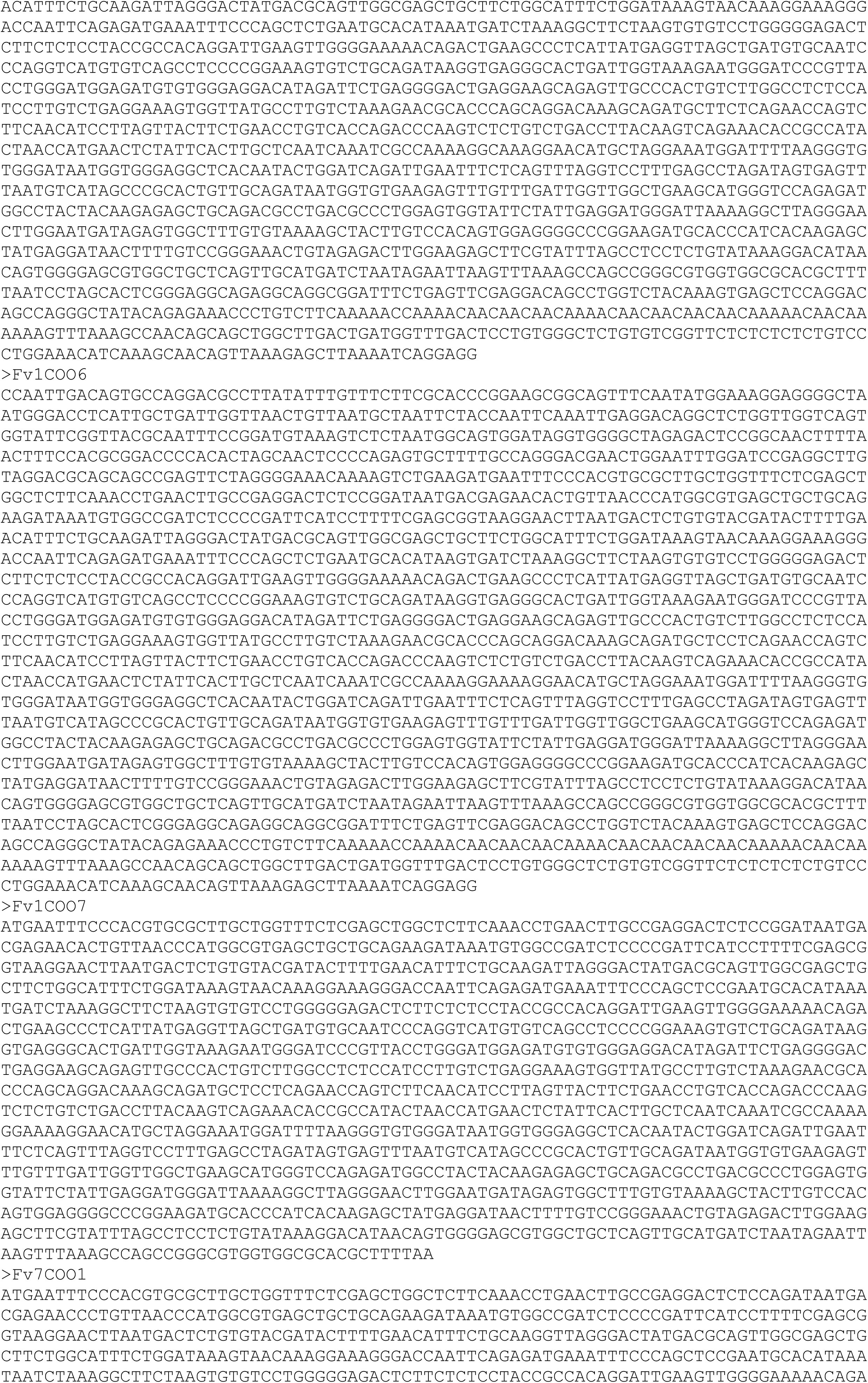
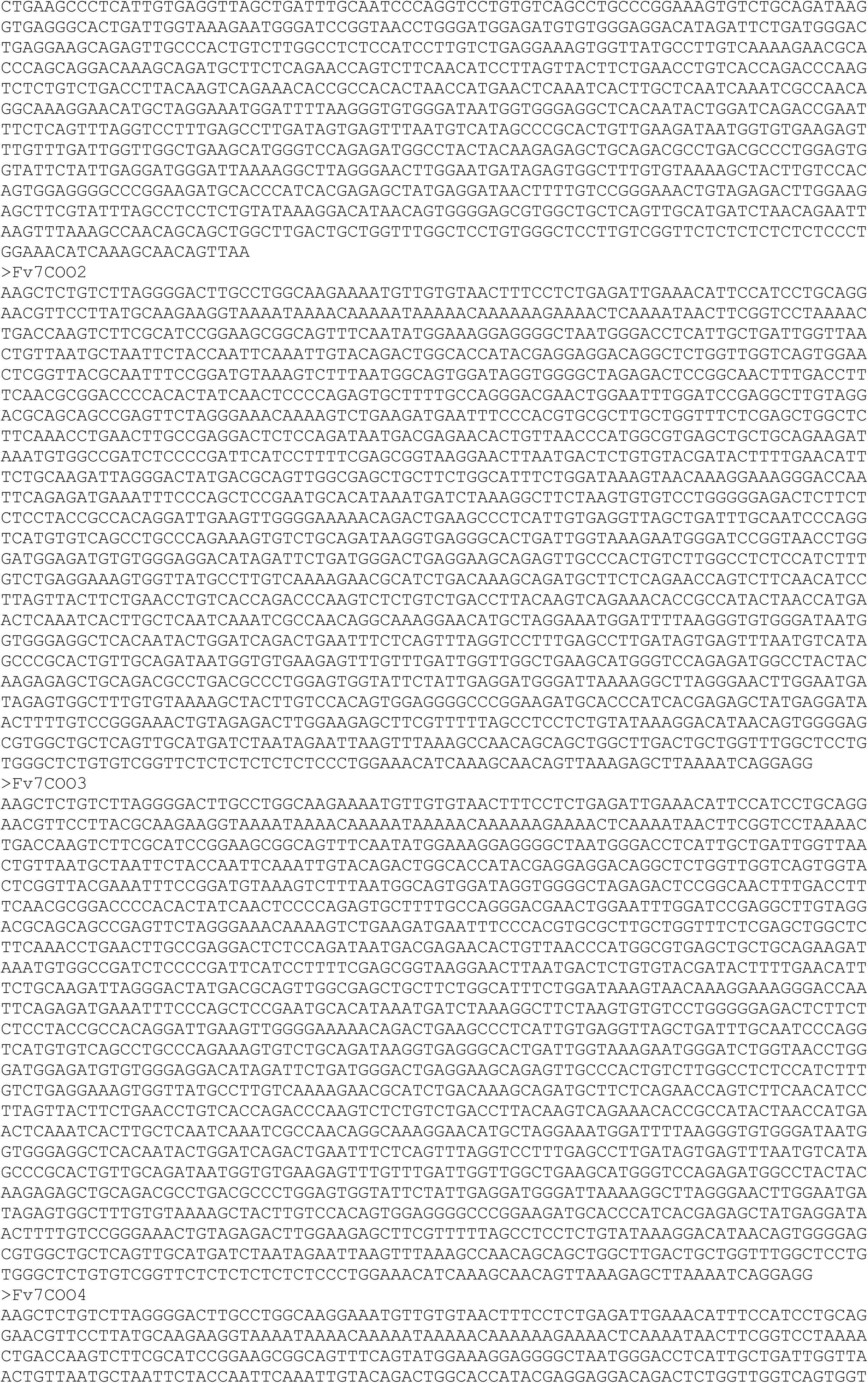
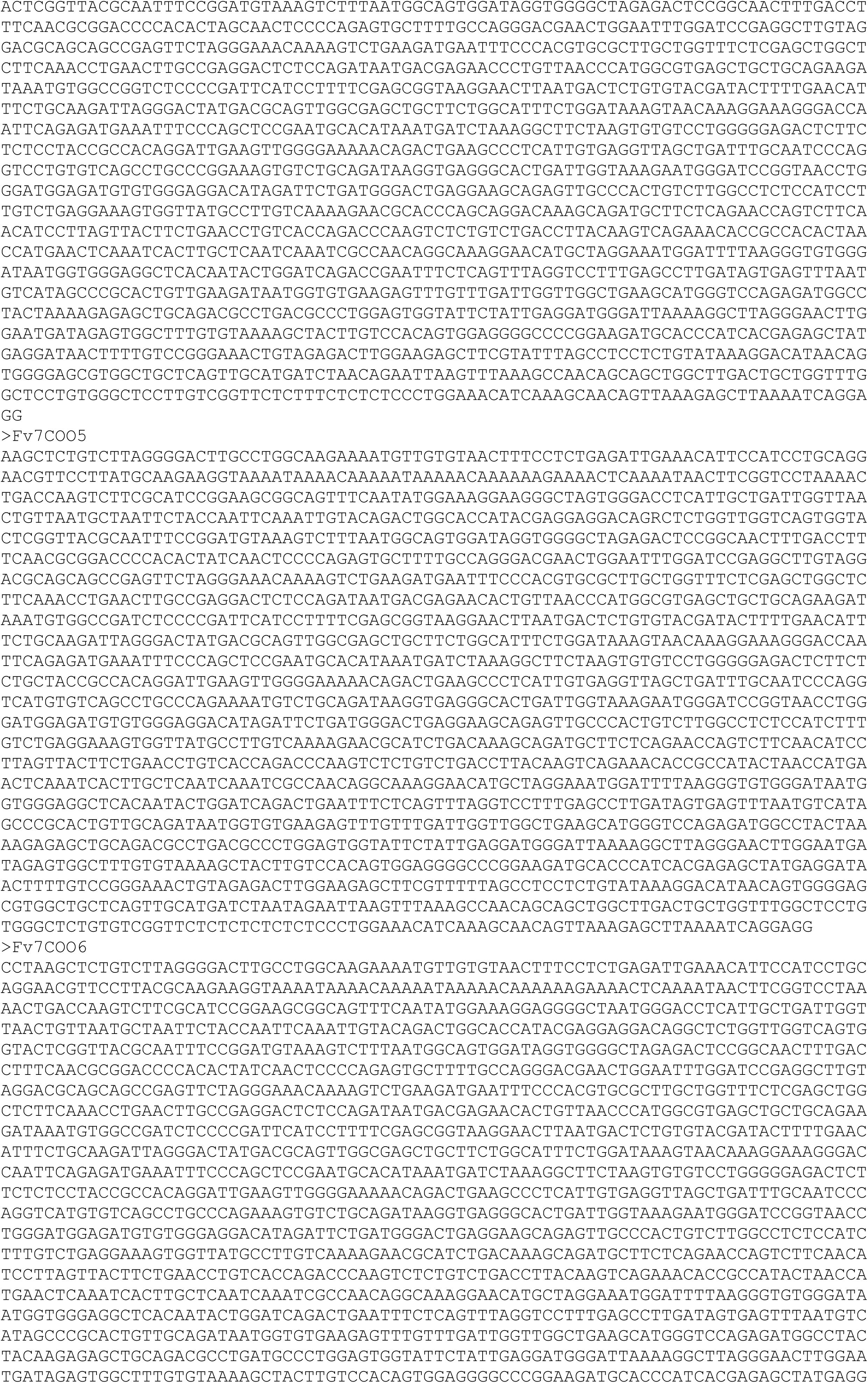
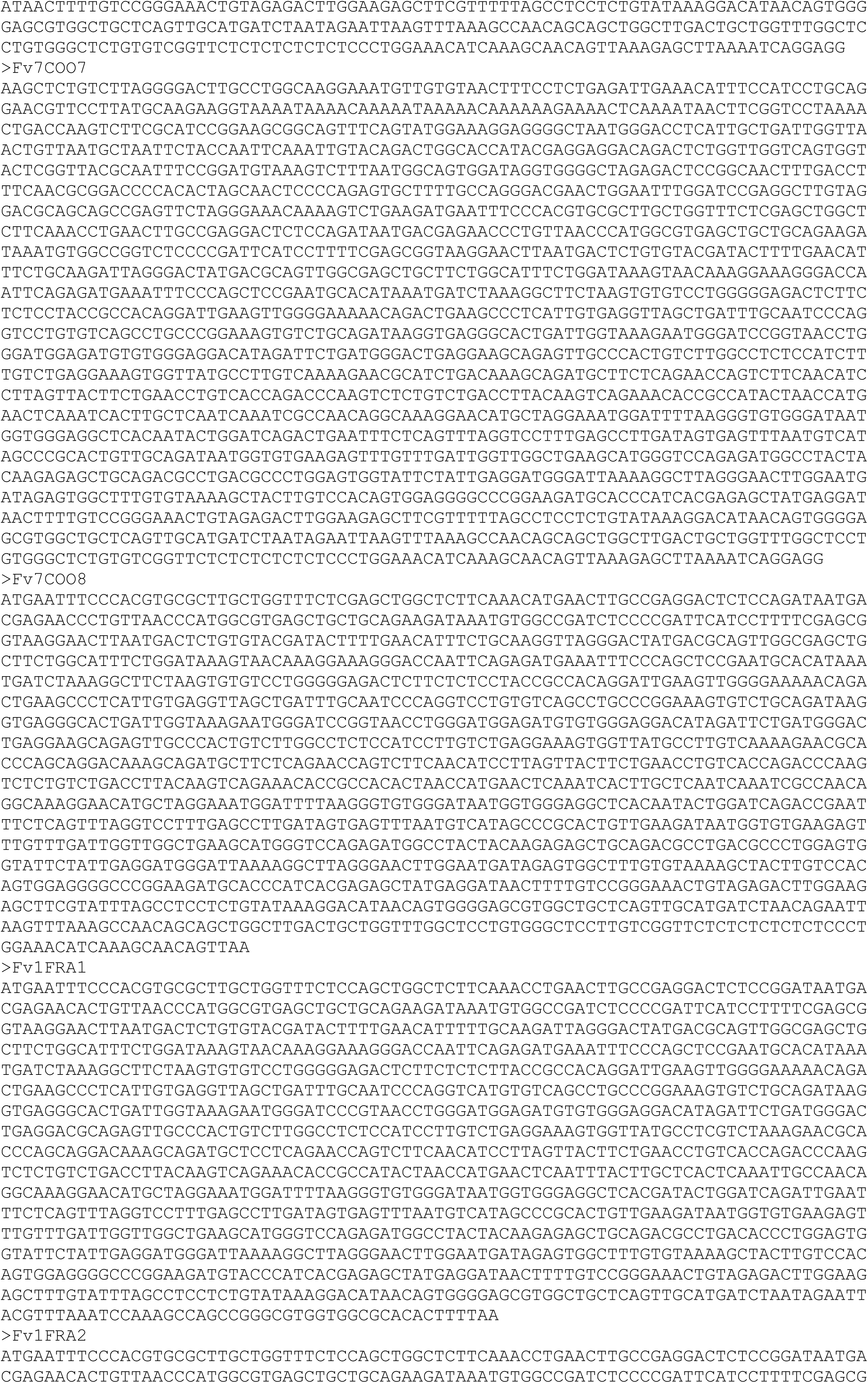
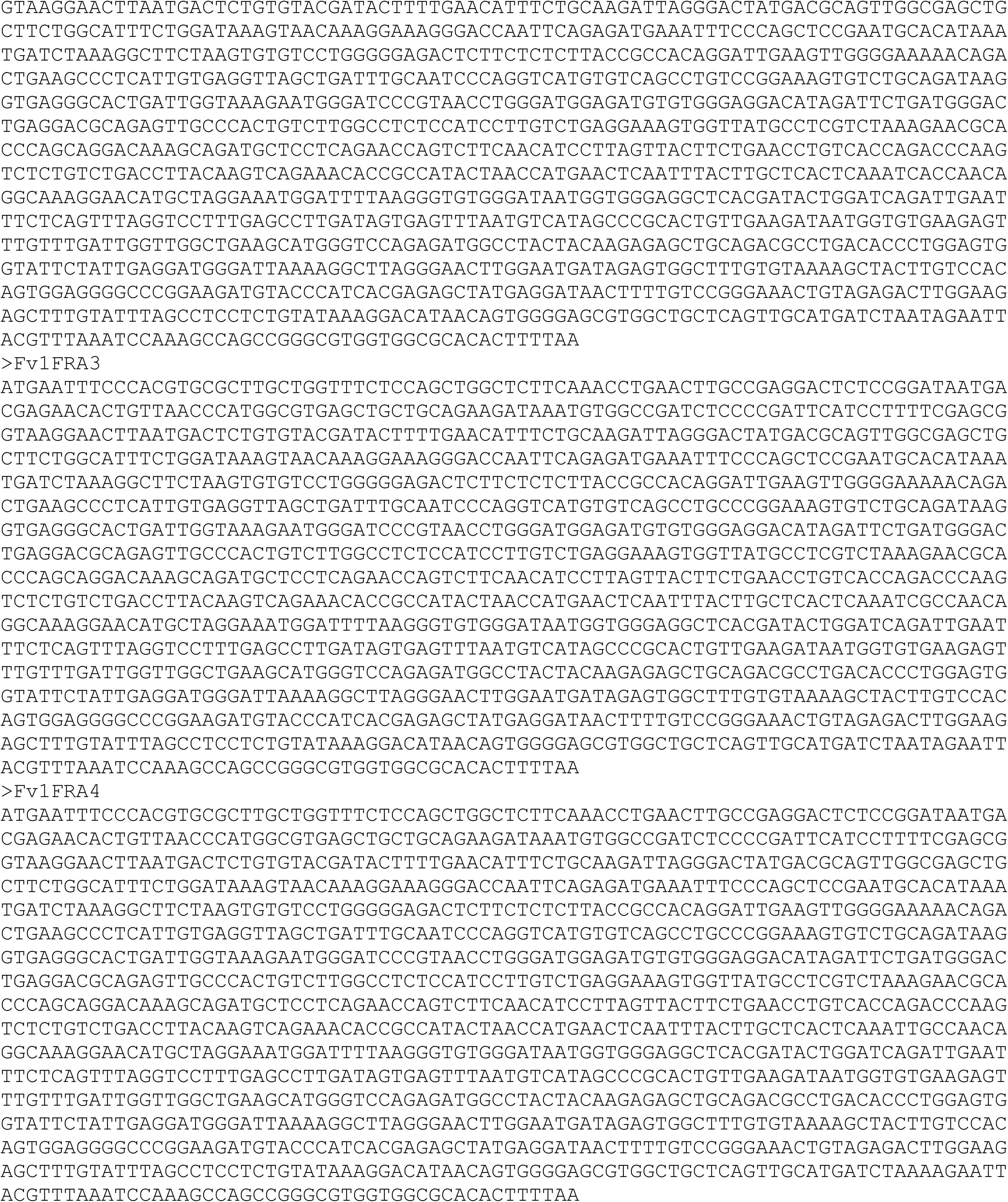
New Fv1 sequences.

